# Bile acid dependent attenuation of toxin mediated disease is independent of colonization resistance against *C. difficile*

**DOI:** 10.64898/2026.03.11.711146

**Authors:** Samantha C. Kisthardt, C. E. Perkins, Abigail S. Gancz, Noah S. Lyons, Stephanie A. Thomas, Emily C. Vincent, Guozhi Zhang, John Tam, Roman Melnyk, Elizabeth C. Rose, Erin S. Baker, Casey M. Theriot

**Affiliations:** Department of Population Health and Pathobiology, College of Veterinary Medicine, North Carolina State University Raleigh, NC 27607; Department of Plant and Microbial Biology, North Carolina State University Raleigh, NC 27607; Center for Gastrointestinal Biology & Disease, UNC School of Medicine, University of North Carolina, Chapel Hill, North Carolina, USA; Department of Chemistry, University of North Carolina at Chapel Hill, Chapel Hill, North Carolina, USA; Molecular Medicine Program, The Hospital for Sick Children Research Institute, 686 Bay Street Toronto, ON, Canada, M5G 0A4; Department of Biochemistry, University of Toronto, Toronto, ON, Canada, M5S 1A8

**Keywords:** *Clostridioides difficile*, *Clostridium scindens*, *Clostridium hiranonis*, bile acids, cholate, deoxycholate, bile salt hydrolase, toxin, inflammation, competition

## Abstract

*Clostridioides difficile* infection (CDI) is a severe antibiotic associated disease and a major cause of morbidity and mortality worldwide. CDI is thought to arise from the loss of protective gut microbes that mediate functions such as secondary bile acid metabolism and nutrient competition, yet the relative contributions of these mechanisms remain unclear. To determine how these processes influence *C. difficile* growth, virulence, and disease, we performed *in vitro* and *in vivo* experiments using two Clostridia strains previously associated with colonization resistance against *C. difficile*. Neither organism prevented colonization or growth through nutrient competition alone. In contrast, secondary bile acid metabolism significantly reduced toxin-mediated disease *in vivo* in a strain dependent manner. These findings demonstrate that secondary bile acid modulation is an important component of CDI prevention independent of nutrient competition and suggest that attenuating virulence, in addition to limiting colonization, may represent a key strategy for next-generation CDI therapeutics.

## INTRODUCTION

*Clostridioides difficile* is an anaerobic, Gram-positive, spore-forming bacterium and a leading cause of diarrheal disease, contributing substantially to morbidity, mortality, and healthcare spending in the United States and worldwide ^1,2^. Antibiotic use increases susceptibility to *C. difficile* infection (CDI) by disrupting the structure and function of the indigenous gut microbiota, thereby creating an environment that supports *C. difficile* colonization ^3–5^. Primary CDI cases are treated with antibiotics such as vancomycin and fidaxomicin, but these treatments can fail and cause up to 25% of patients to develop recurrent CDI (rCDI) ^6–9^. Following an initial recurrence, patients are at substantially higher risk for multiple, more severe episodes of rCDI ^10,11^. One strategy to break the rCDI cycle and prevent future episodes is fecal microbiota transplantation (FMT), which works by recolonizing the recipient with healthy donor microbiota ^12^. While highly effective, instances of FMT-associated antimicrobial resistance transfer and other adverse outcomes have driven the development of FDA-approved alternative treatments, herein called microbiota focused therapeutics (MFTs) ^13–15^. Although similarly effective to FMT, MFTs are cost-prohibitive, exclusively available in the United States, not authorized for pediatric patients or for fulminant CDI, and remain donor dependent ^16,17^. Consequently, there is an urgent need to develop new, non-donor derived therapies. These emerging therapeutics, termed live biotherapeutic products (LBPs), consist of well-defined consortia of live bacteria ^18,19^. While LBPs such as VE303, a consortium of eight bacterial strains, are currently in clinical trials for rCDI, none are FDA-approved ^20^. One of the challenges of rationally developing these consortia is a limited mechanistic understanding of how specific bacteria, alone and in tandem, can prevent *C. difficile* colonization and virulence.

The ability of the gut microbiota to prevent intestinal colonization by pathogens, termed colonization resistance (CR), is mediated by multiple mechanisms ^21^. In the case of *C. difficile*, CR has historically been associated with the presence of microbially-produced secondary bile acids (BAs), which inhibit multiple stages of the *C. difficile* lifecycle, and the presence of commensal Clostridia like Lachnospiraceae ^3,4,22–24^. However, the mechanism of protection derived from these BAs and microbes is still unclear, particularly because most studies have been associative. While primary BAs are synthesized by the host from cholesterol and conjugated to either glycine or taurine ^25^, secondary BA production requires bacteria encoding bile salt hydrolases (BSHs) to first deconjugate the host-conjugated amino acid from the primary sterol core to produce the unconjugated primary BAs cholate (CA) or chenodeoxycholate (CDCA) ^25,26^. Only after BSH-catalyzed deconjugation can a relatively small number of bacteria, mainly Clostridia, transform primary BAs to secondary BAs. This is performed via the *bai* operon, which consists of eight enzymes that collectively perform the multistep process of 7[7]-dehydroxylation to yield deoxycholate (DCA) from CA or lithocholate (LCA) from CDCA ^25,27^. Of the *bai* encoding bacteria, *Clostridium scindens* is the most well studied, however mouse models with *C. scindens* and *C. difficile* have shown inconsistent results in restoring CR against *C. difficile*, requiring further scrutiny ^22,23,28,29^. Recently, it was also demonstrated that some BSHs are able to further modify both primary and secondary BAs by reconjugating non-canonical amino acids onto BA cores, forming microbially-conjugated BAs (MCBAs) that can differentially alter the *C. difficile* lifecycle *in vitro and in vivo* ^24,30,31^. This finding has significantly increased the complexity of the BA pool in the gut, requiring new scrutiny when defining the relationship between these BAs and *C. difficile*.

Despite the evidence for the role of BAs in CDI-associated CR, recent work concludes that CR against *C. difficile* is BA-independent. Instead, these studies suggest CR is solely driven by competition for nutrients, specifically amino acids such as proline and glycine, which are essential for Stickland fermentation, a metabolic process unique to Clostridia ^32^. Both bioreactor and mouse models show that competition for these nutrients is associated with protection against *C. difficile* ^23,33–35^. Another study showed that the consumption of Stickland amino acids by Peptostreptococcaceae *(e.g., Paraclostridium bifermentans*, *Clostridium hiranonis*) could alter the host nutrient landscape in a gnotobiotic mouse model, causing *C. difficile* to use less favorable metabolic pathways and leading to decreased growth and virulence ^33,36^. One interpretation of these studies is that competition for nutrients by commensal Clostridia is sufficient for providing CR against *C. difficile*, while secondary BA metabolism is dispensable for protection.

Based on these recent findings and prior work on the importance of BAs, we hypothesize that nutrient competition, in addition to Clostridia-driven modifications of the BA pool, confer CR against *C. difficile*. To test the interplay between nutrient competition and BA modifications, we selected two gut-associated commensal Clostridia species, the non-antimicrobial peptide encoding *Clostridium scindens* VPI12708 and *Clostridium hiranonis* TO-931. While both organisms use Stickland fermentation, produce secondary BAs via the *bai* operon, and further modify BAs via 7[7]- and 12[7]-hydroxysteroid dehydrogenases (HSDHs), *C. hiranonis* is also predicted to encode a BSH ^37–40^. Taking advantage of the differential BA profiles produced by these Clostridia strains due to differences in their encoded BA-modifying enzymes, we leveraged *in vitro* and *in vivo* models in addition to metabolomic and transcriptomic approaches to investigate the extent to which CR against *C. difficile* is BA dependent or independent.

Here we show that neither *C. scindens* nor *C. hiranonis* alone were able to prevent *C. difficile* growth *in vitro* or colonization *in vivo*, suggesting that competition for nutrients alone with these bacteria does not provide CR or protection from *C. difficile*. Although neither strain was able to decrease *C. difficile* colonization in a gnotobiotic mouse model, C*. hiranonis* was able to suppress *C. difficile* virulence and attenuate toxin mediated disease. Complementary *in vitro* assays only mirrored the toxin attenuation by suppression of activity seen *in vivo* when the primary BA cholate was present, suggesting this phenotype is BA dependent. To identify the mechanism of this protection, we leveraged targeted BA metabolomics and transcriptomics on all *in vitro* and *in vivo* samples with this shared phenotype. Subinhibitory concentrations of secondary bile acid DCA made by *C. hiranonis* in the presence of CA were responsible for suppressed virulence by *C. difficile*. Additionally, we show that the BSH encoded by *C. hiranonis*, but absent in *C. scindens*, is important for DCA mediated protection *in vivo*. This previously undescribed role of DCA in protecting against toxin-mediated disease independently of CR highlights that the physiological functions of BAs are more nuanced than previously appreciated. It is critical to define how these bacteria and their metabolic products impact both CR and virulence factors, which will inform the rational design of targeted bacterial therapeutics for the treatment of CDI and other intestinal diseases beyond CDI.

## RESULTS

### *C. scindens* affects *C. difficile* growth and toxin activity in a BA concentration-dependent manner in the absence of competitive pressure *in vitro*

We first sought to determine if there is competition between *C. scindens* and *C. difficile in vitro* both in the absence and presence of CA, which should be converted into DCA by the *C. scindens bai* operon ^41^. We chose to use a subinhibitory concentration of CA (1.25 mM) because higher concentrations (2.5 mM CA) affected commensal growth in coculture. *C. scindens* does not decrease *C. difficile* growth or toxin activity in coculture, regardless of whether 1.25 mM CA is supplemented, as determined by comparison to the respective *C. difficile* monoculture with or without CA (Figure 1A and C). There was also no difference in *C. difficile* growth in monoculture in the presence of CA compared to those in coculture (Welch’s t-test p=0.0117). In contrast, *C. scindens* growth decreases in coculture when CA is present (Welch’s t-test p=0.0008), suggesting that BA metabolism may negatively impact *C. scindens* fitness in the presence of competitive pressure (Figure 1A). Adding DCA, which should mimic *C. scindens’* ability to convert CA to DCA, again decreased *C. difficile* growth in the monoculture, but the levels were similar in the coculture. This lower *C. difficile* load in the presence of DCA corresponded to decreased toxin activity, which is expected as growth and toxin are interdependent (Figure 1A, Welch’s t-test p=0.0332 and Figure 1C p=0.0006). *C. scindens* growth was restored in coculture with DCA (Figure 1A), suggesting that DCA alone is not driving the inhibition of *C. scindens*, and that decreased growth in coculture with *C. difficile* and CA could be due to the metabolic needs associated with the production of DCA by the *bai* operon. This shows that 1.25 mM CA negatively influences *C. scindens*’ ability to compete with *C. difficile in vitro* but does not affect *C. difficile* growth.

**Figure 1.**
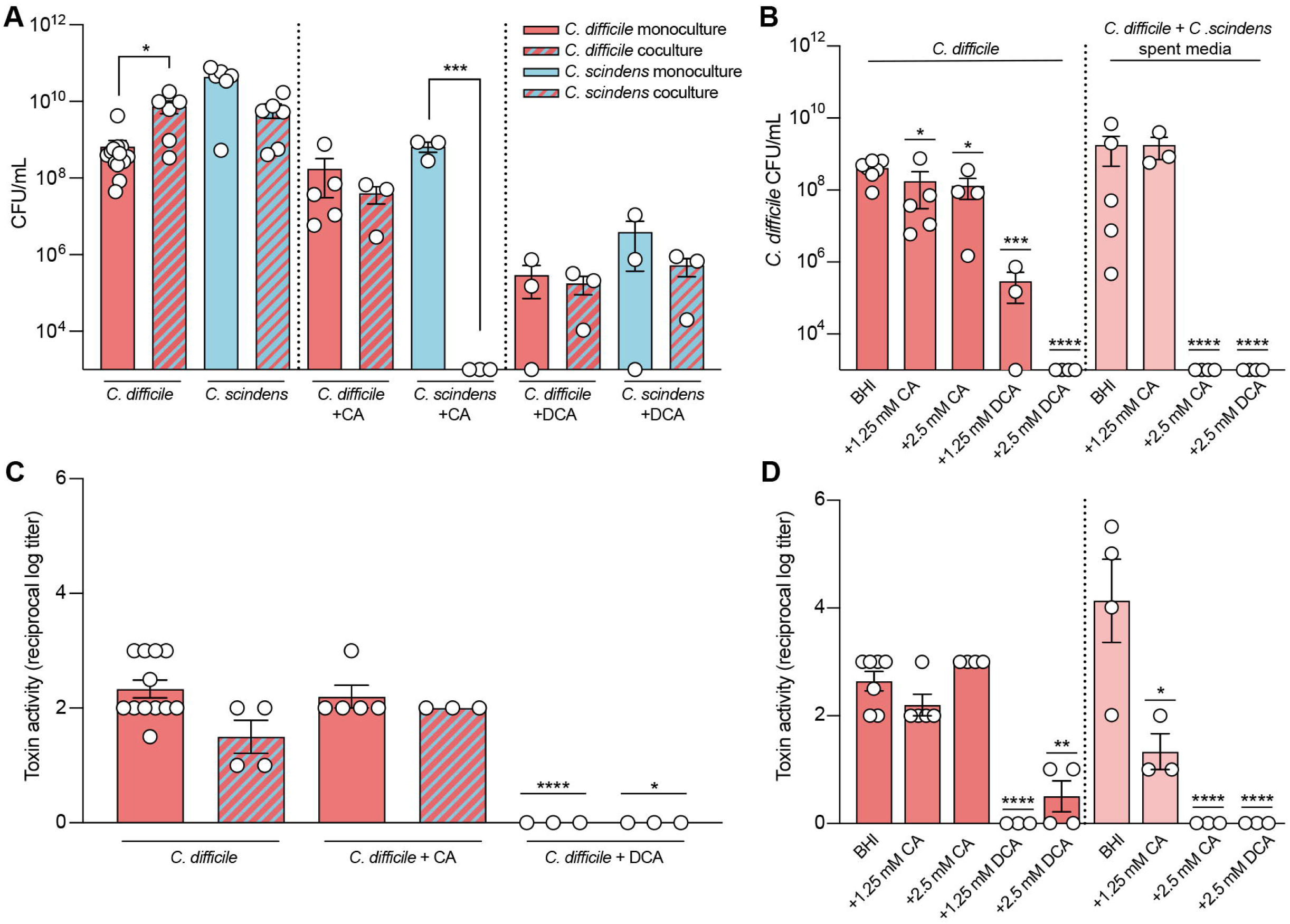
*C. scindens* affects *C. difficile* growth and toxin activity in a BA concentration-dependent manner in the absence of competitive pressure *in vitro*. A) Competition study to measure growth of *C. difficile* and *C. scindens* in monoculture (solid bars) or 1:1 coculture (striped bars) without BAs or supplemented with either 1.25 mM CA or 1.25 mM DCA after 24 hr of incubation. Statistical comparisons were made between monoculture and coculture conditions for each organism by Welch’s t-test. B) Spent media study to measure *C. difficile* growth at 24 hr after supplementation with 1.25 mM (low) and 2.5 mM (high) CA or DCA or filtered spent media from cultures of C. *scindens* grown with low and high CA or DCA. Statistical comparisons were made between *C. difficile* cultures and their respective *C. scindens* spent media culture by Welch’s t-test. C) Toxin activity of cultures in (A). Activity was quantified by Vero cell cytotoxicity assay and statistical comparisons were made between *C. difficile* monoculture and coculture for each BA condition by Welch’s t-test. D) Toxin activity of cultures in (B). Activity was quantified by Vero cell cytotoxicity assay and statistical comparisons were made between *C. difficile* cultures and their respective *C. scindens* spent media culture by Welch’s t-test. For figures (A-D), mean of biological replicates (n = 3-10) is graphed and error bars represent the SEM. * p < 0.05, ** p < 0.01, *** p < 0.001, **** p < 0.0001.

We next grew *C. difficile* in spent media from *C. scindens* grown alone or in the presence of 1.25 mM (low) or 2.5 mM (high) CA to determine if bacterial derived products, including secondary BAs from *C. scindens*, can alter *C. difficile* growth and virulence in the absence of competition. While *C. scindens* spent media did not alter *C. difficile* growth or toxin activity (Figure 1B and D), spent media from *C. scindens* supplemented with low CA was able to decrease toxin activity, but not growth (Figure 1B and D, Welch’s t-test p = 0.0361). However, spent media from *C. scindens* supplemented with high CA inhibited both *C. difficile* growth and toxin activity (Figure 1B and D, Welch’s t-test p<0.0001). We next looked at *C. difficile* cultures supplemented with CA or DCA (low and high) to determine whether these observed effects are due to *C. scindens*-mediated BA metabolism. CA alone (low and high) had slight effects on *C. difficile* growth (Figure 1B Welch’s t-test P = 0.02) but did not affect toxin activity, while the addition of DCA (low and high) decreased growth and toxin activity (Figure 1B and D Welch’s t-test p = 0.0001, p<0.0001). This aligns with previous findings, where *C. scindens* could produce roughly 2 mM DCA in the presence of 2.5 mM CA ^41^. These trends were replicated in *C. difficile* cultures grown in spent media from *C. scindens* high DCA (Figure 1B and D, Welch’s t-test p<0.0001), indicating that the presence of DCA was responsible for the effects on *C. difficile*. *C. scindens* alone cannot reduce *C. difficile* growth and virulence, it requires CA to make DCA only when it is not in direct competition with *C. difficile*. *C. scindens* affects *C. difficile* growth and virulence differently in a BA concentration dependent manner.

### *C. scindens* does not prevent *C. difficile* colonization or disease *in vivo*

To test whether *C. scindens* can inhibit *C. difficile* colonization or disease in a host, which has a more complex BA pool, we used a gnotobiotic mouse model of CDI. Germ-free (GF) mice either remained GF or were challenged with *C. scindens* and monocolonization was monitored over a seven-day period, or days -7 to day 0 post *C. difficile* challenge (Figure 2A, Figure S1A-B). On day 0, mice were challenged with *C. difficile* spores (Figure 2A). After two days, mice monocolonized with *C. scindens* did not lose weight, whereas mice monocolonized with *C. difficile* or cocolonized with *C. scindens* and *C. difficile* lost weight (Figure 2B, Kruskal-Wallis test with Dunn’s correction p=0.0015). *C. difficile* load was similar between these groups on day 2 post *C. difficile* challenge, while *C. scindens* load was higher in cocolonized mice (Figure 2C, Mann-Whitney test p= 0.0182). *C. difficile* toxin activity was slightly decreased in cocolonized mice when compared to *C. difficile* monocolonized mice (Figure 2D, Mann-Whitney test p=0.0205) and spore load did not change across groups (Figure 2E). Cocolonized mice had inflammation and epithelial damage similar to *C. difficile* monocolonized mice, while edema was lower (Figure 2F and G, Two-way ANOVA with Tukey’s multiple comparisons test p=0.0062). There were no differences in colonic histopathology between *C. difficile* monocolonized mice and cocolonized mice (Figure S1C-D Two-Way ANOVA with Tukey’s Multiple Comparison Test). These results demonstrate that, as in our *in vitro* results, *C. scindens* cannot prevent colonization or reduce the virulence of *C. difficile* via nutrient competition alone *in vivo*.

**Figure 2.**
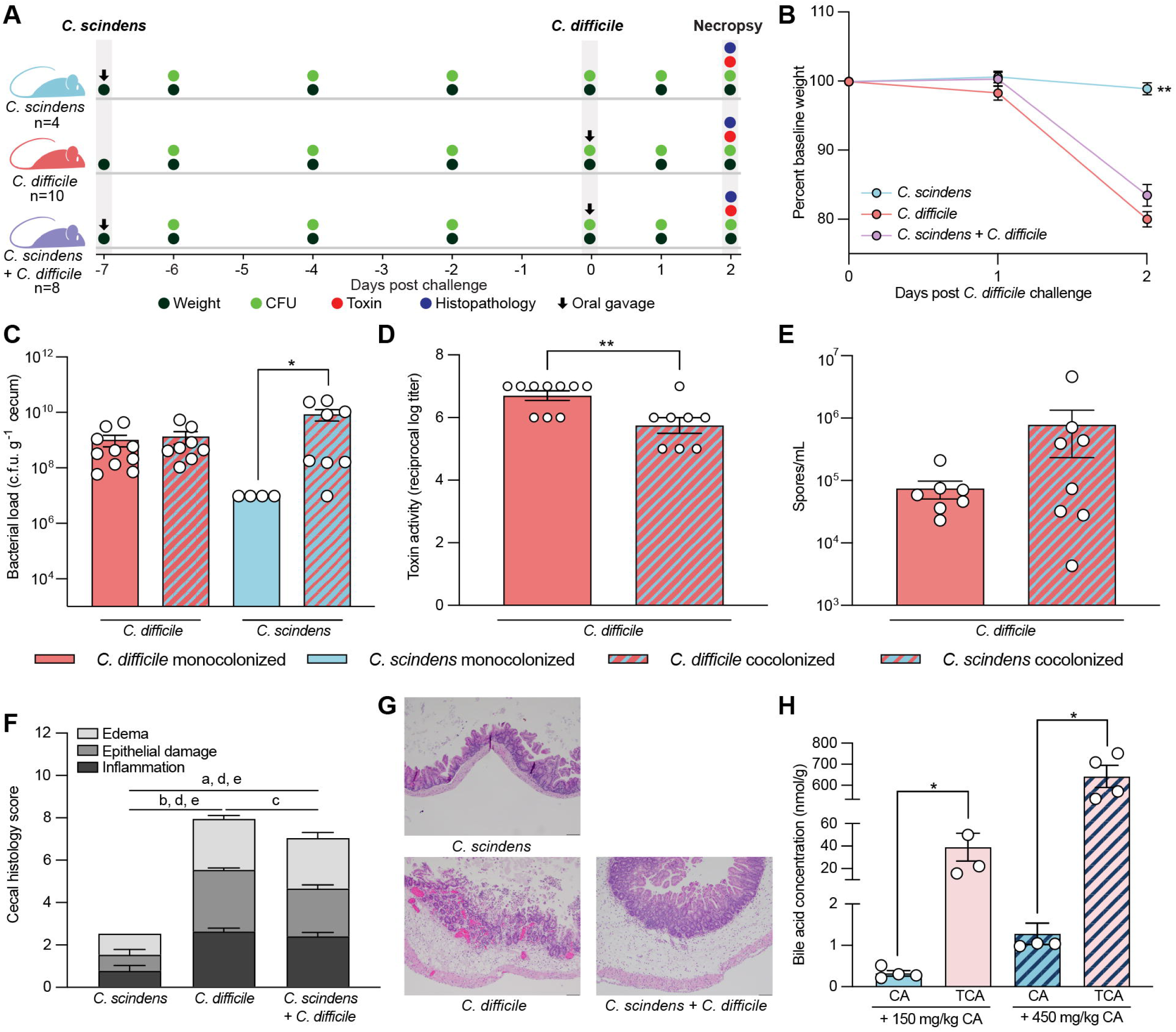
*C. scindens* does not prevent *C. difficile* colonization and disease *in vivo*. A) Experimental overview of the gnotobiotic mouse model. B) Percent change in weight from baseline was tracked from days 0-2 post-*C. difficile* challenge in *C. scindens* monocolonized mice (n = 4), *C. difficile* monocolonized mice (n = 10), and cocolonized mice (n = 8). Statistical significance was determined by using Kruskal-Wallis test with Dunn’s correction to compare each condition to the *C. difficile* monocolonized mice. C) *C. difficile* bacterial load in cecal content collected at day 2. Statistical comparisons were made between monocolonized (solid bars) and cocolonized (striped bars). D) *C. difficile* toxin activity measured by Vero cell cytotoxicity assay on cecal content from (C). Statistical significance was determined by Mann-Whitney test, comparing *C. difficile* monocolonized mice to cocolonized mice. E) *C. difficile* spore load from (C). F) Histopathological changes to cecal tissue from each treatment. Tissues were scored for edema, epithelial damage, and inflammation on a scale of 0-4 for a maximum total score of 12 and statistical comparisons were made between each scoring category across all conditions by two-way ANOVA (a, p < 0.01 edema; b, p < 0.001 edema; c, p < 0.05 epithelial damage; d, p < 0.01 epithelial damage; e, p < 0.01 inflammation). G) Representative photomicrographs of tissue collected at necropsy from (F). H) Concentration of CA (blue) and TCA (pink) in cecal content of mice given 150 mg/kg CA (solid bars) or 450 mg/kg CA (striped bars) in drinking water. Significance was determined by comparing the amount of CA and TCA present in each condition. For figure (B) each point represents the mean of all biological replicates for each treatment, and error bars represent SEM. Figures (C-F and H) show each bar representing the mean of all biological replicates for each treatment, and error bars represent the SEM. * p < 0.05, ** p < 0.01.

To determine if *C. scindens* can make the secondary BA DCA *in vivo*, we performed targeted BA metabolomics on cecal content collected at day 2. While *C. scindens* produces DCA *in vitro*, we did not detect DCA *in vivo* (Figure S1E) ^41^. The lack of DCA *in vivo* could be due to the lack of free CA, as it is in the form of host conjugated TCA that must be deconjugated by a BSH to become accessible ^42^. Finally, when comparing the BA pools of *C. difficile* monocolonized mice to those of GF mice there was an increase in 7-oxoDCA, which is produced by 7[7]HSDH activity, alongside the primary BAs CA, CDCA, MCA, UDCA, and CA (Figure S1F, t-test). The increase in 7-oxoDCA suggests that *C. difficile* can modify CA into oxo-bile acids *in vivo*, which is supported by a study showing *C. difficile* encodes a 7[7]HSDH ^33,43^.

We next tested whether, as it did *in vitro*, supplementing excess CA could help protect mice against CDI. Mice were given *C. scindens* and CA was supplemented in their drinking water (150 mg/kg), starting on day -2, or two days prior to *C. difficile* challenge. Supplementing CA in the drinking water of mice monocolonized with *C. scindens* did not provide protection against CDI, as the results were similar to those of the no CA condition (Figure S1G-I). We next leveraged BA metabolomics to determine the amount of available CA in GF mice given a low amount of CA (150 mg/kg) and a high amount of CA (450 mg/kg) in their drinking water. While CA did increase in the mice, TCA also increased in a dose-dependent manner, suggesting that most of the supplemented CA could be host-conjugated with taurine and inaccessible to *C. scindens*, as it lacks a BSH, which is required to deconjugate TCA into free CA (Figure 2H, Mann-Whitney test p = 0.0286).

### *C. hiranonis* does not inhibit *C. difficile* growth but does inhibit toxin activity in a BA concentration dependent manner *in vitro*

Next, we wanted to compare how another Clostridia with different BA altering enzymes, *C. hiranonis*, which encodes a BSH in addition to the *bai* operon, would affect *C. difficile* growth and toxin activity *in vitro*. As with *C. scindens*, *C. hiranonis* does not alter *C. difficile* growth or toxin activity in coculture in the absence or presence of 1.25. mM CA (Figure 3A and C). In contrast, *C. hiranonis* growth is inhibited in coculture with *C. difficile* compared to its respective monoculture (Figure 3A, Welch’s t-test p = 0.0004), which aligns with previous work by our group ^41,44^. However, the presence of CA rescues *C. hiranonis* growth in coculture (Figure 3A). The addition of DCA further decreased *C. hiranonis* growth in both cultures, suggesting that the rescue effect of CA supplementation may not be due to the production of DCA alone (Figure 3A). These data indicate that *C. hiranonis* cannot inhibit *C. difficile*, however supplementation of CA rescues *C. hiranonis* growth in direct competition with *C. difficile*.

**Figure 3.**
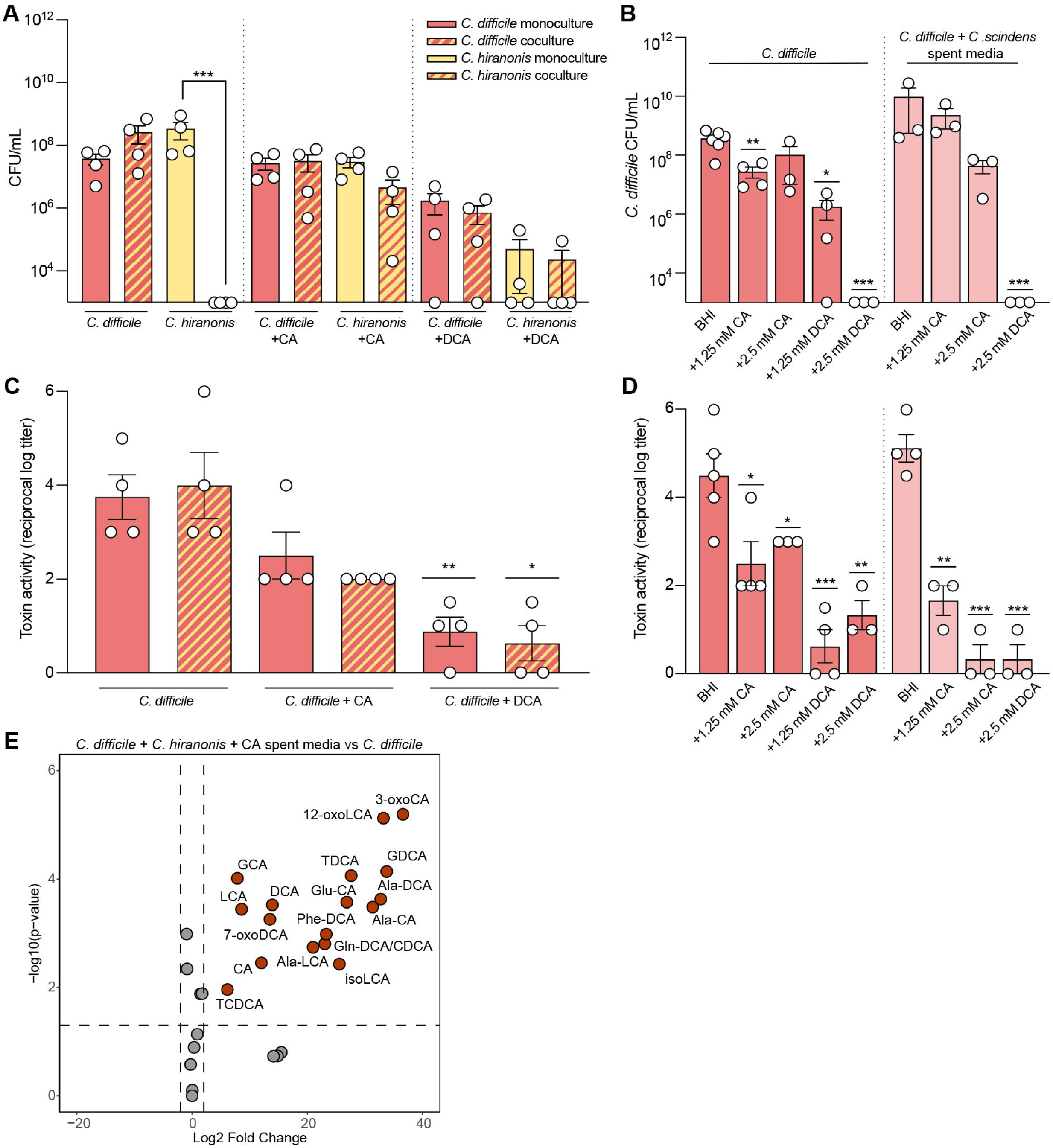
*C. hiranonis* supplemented with cholate decreases *C. difficile* toxin activity in a growth-independent manner *in vitro*. A) Growth of *C. difficile* and *C. hiranonis* monoculture (solid bars) or 1:1 coculture (striped bars) without BAs or supplemented with either 1.25 mM CA or 1.25 mM DCA after 24 hr. Statistical comparisons were made between monoculture and coculture conditions for each organism by Welch’s t-test. (B) *C. difficile* growth at 24 hr in fresh BHI supplemented with 1.25 mM (low) and 2.5 mM (high) CA or DCA or filtered spent media from cultures of C. *hiranonis* grown with 1.25 mM (low) and 2.5 mM (high) CA or 2.5 mM (high) DCA. Statistical comparisons were made between *C. difficile* cultures in fresh BHI and their respective *C. hiranonis* spent media culture by Welch’s t-test. (C) Toxin activity of cultures in (A). Activity was quantified by Vero cell cytotoxicity assay and statistical comparisons were made between *C. difficile* monoculture and coculture for each bile acid condition by Welch’s t-test. (D) Toxin activity of cultures in (B). Activity was quantified by Vero cell cytotoxicity assay and statistical comparisons were made between *C. difficile* cultures in fresh BHI and their respective *C. hiranonis* spent media culture by Welch’s t-test. (E) Volcano plot of bile acids detected in cultures of *C. difficile* grown in spent media from *C. hiranonis* +/- 2.5 mM CA compared to cultures of *C. difficile* alone. Statistical significance was determined by Student’s t-test (log2 fold change > 2, adjusted p value < 0.05). For figures (A, C) bars represent the mean of all biological replicates for each treatment (n = 4-8). For figures (B, D) bars represent the mean of all biological replicates for each treatment (n = 3-8). All error bars represent the SEM. Figure (E) was generated by comparing n = 3 biological replicates from each condition. * p < 0.05, ** p < 0.01, *** p < 0.001.

Since the addition of CA drives *C. hiranonis* growth in the presence of *C. difficile*, we wanted to determine whether *C. hiranonis* might be able to modify 1.25 mM (low) or 2.5 mM (high) CA to produce bacterial products that could affect *C. difficile* growth or toxin activity in the absence of competition. *C*. *hiranonis* spent media alone did not alter *C. difficile* growth or toxin activity (Figure 3B and D). Similarly, while spent media from *C. hiranonis* with low and high CA did not affect *C. difficile* growth, toxin activity decreased in both CA conditions (Figure 3B and D, Welch’s t-test p = 0.0053 and p = 0.0003). The addition of *C. hiranonis* spent media with high DCA decreased *C. difficile* growth and toxin activity (Figure 3B and 3D, Welch’s t-test p < 0.0001, p = 0.0003). We next supplemented cultures of *C. difficile* with low and high CA or DCA to test whether these effects are BA mediated. Low CA caused a slight decrease in growth (Figure 3B Welch’s t-test p = 0.0041), while low and high CA decreased toxin activity slightly (Figure 3D Welch’s t-test p = 0.0379, p = 0.0476). In comparison, low and high DCA decreased growth (Figure 3B Welch’s t-test p = 0.029, p < 0.0001) and decreased toxin activity at or near the limit of detection (Figure 3D Welch’s t-test p = 0.0003, p = 0.0016). These results suggest that while *C. hiranonis* produces less DCA than *C. scindens,* it can decrease *C. difficile* toxin activity in a growth-independent manner at both low and high concentrations of CA ^41^.

We next wanted to define the BAs in spent media that suppressed *C. difficile* toxin activity in a growth independent manner. We measured the BAs in *C. difficile* cultures grown in spent media from *C. hiranonis* supplemented with high CA and compared the resulting BA pool to that of *C. difficile* grown alone. Many BAs increased, including the secondary BAs DCA and LCA, and the canonical conjugated BAs GCA, TCDCA, TDCA, and TDCA (Figure 3E, t-test). Products of HDSH activity like 3-oxoCA, 7-oxoDCA, 12-oxoLCA, and isoLCA were also increased, as were the BSH-produced MCBAs Ala-CA, Glu-CA, Ala-DCA, Gln-DCA/CDCA, and Phe-DCA (Figure 3E, t-test). This suggests that one of these BAs alone, or several BAs in combination, could be responsible for the suppression to *C. difficile* toxin activity independent of growth.

### *C. hiranonis* does not prevent *C. difficile* colonization but does ameliorate disease *in vivo*

Next, we wanted to determine if *C. hiranonis* was able to alter *C. difficile* growth and virulence in a gnotobiotic mouse model of CDI. As with previous experiments, we monocolonized mice with *C. hiranonis* for seven days and monitored both colonization and weight loss before challenging with *C. difficile* spores on day 0 (Figure 4A, Figure S2A-B). Mice cocolonized with *C. hiranonis* and *C. difficile* lost less weight than *C. difficile* monocolonized mice (Figure 4B, Kruskal-Wallis test p = 0.0005). While we did not observe any differences in *C. difficile* or *C. hiranonis* load across mono- or cocolonized mice (Figure 4C), we found that *C. difficile* toxin activity and spore load was lower in cocolonized mice (Figure 4D, Mann-Whitney test p = 0.0010) (Figure 4E, Welch’s t-test p = 0.0012). Cocolonized mice also had less edema, epithelial damage, and inflammation than *C. difficile* monocolonized mice (Figure 4F and G, Two-way ANOVA with Tukey’s Multiple Comparisons Test edema p = 0.0102, epithelial damage p = 0.0002, inflammation p = 0.0003). But this protection was not seen in the colon (Figure S2C-D). Subsequently, we also wanted to determine whether CA supplementation could enhance protection against CDI. Supplementing CA (150 mg/kg) in the drinking water of mice monocolonized with *C. hiranonis* did not provide additional protection against CDI, as the results were similar to the no CA condition (Figure S2E-G). Regardless of CA supplementation, *C. hiranonis* challenge can prevent *C. difficile* toxin-mediated host tissue damage in a growth-independent manner. Next, we wanted to determine whether *C. hiranonis*-mediated protection against *C. difficile* toxin mediated disease was short lived, or whether it could be sustained longer than two days. To do so, we extended the model to four days post challenge. In this extended model, weight loss, bacterial load, toxin activity, and histopathological changes were similar to day 2, suggesting that *C. hiranonis* can provide sustained protection against CDI (Figure 2 SH-L).

**Figure 4.**
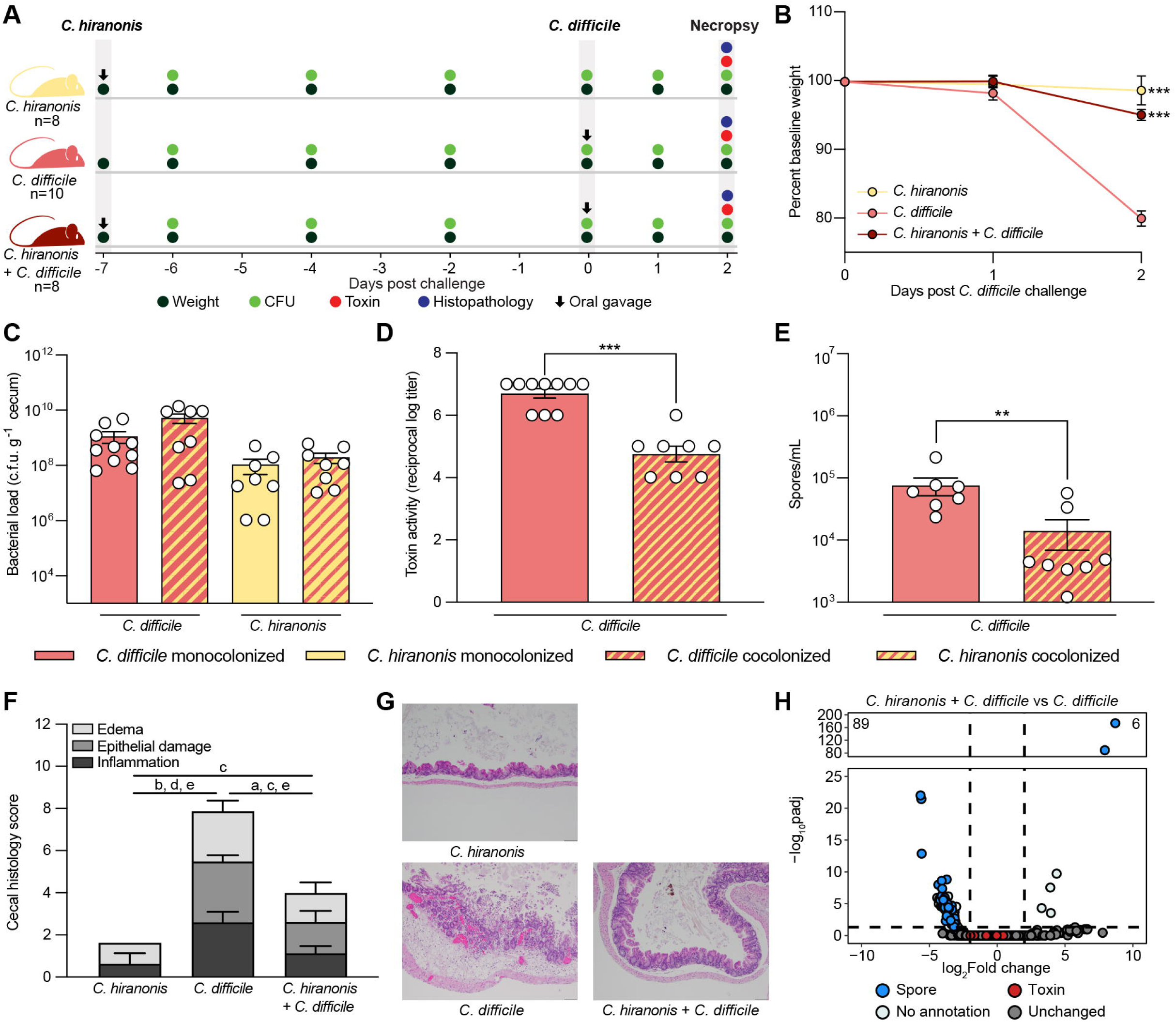
*C. hiranonis* does not prevent *C. difficile* colonization, but ameliorates disease *in vivo*. A) Experimental overview of the gnotobiotic mouse model. B) Percent change in weight from baseline from days 0-2 post-*C. difficile* challenge in *C. hiranonis* monocolonized mice (n = 8), *C. difficile* monocolonized mice (n = 10), and cocolonized mice (n = 8). Statistical significance was determined by using Kruskal-Wallis test with Dunn’s correction to compare each condition to the *C. difficile* monocolonized mice. C) *C. difficile* bacterial load in cecal content collected at day 2. D) *C. difficile* toxin activity measured by Vero cell cytotoxicity assay on cecal contents from (C). Statistical significance was determined by comparing *C. difficile* monocolonized mice to cocolonized mice with a Mann-Whitney test. E) *C. difficile* spore load from (C). Statistical significance was determined by Welch’s t-test. F) Histopathology changes to cecal tissue from each treatment. Tissues were scored for edema, epithelial damage, and inflammation on a scale of 0-4 for a maximum total score of 12 and statistical comparisons were made between each scoring category across all conditions by two-way ANOVA (a, p< 0.01 edema; b, p<0.0001 edema; c, p<0.01 epithelial damage; d, p<0.0001 epithelial damage; e, p< 0.0001 edema). G) Representative photomicrographs of tissue collected from mice in F. H) Volcano plot of RNAseq DESeq2 output showing changes to *C. difficile* gene expression in cocolonized mice compared to monocolonized mice. Significant differential expression was defined as any gene with +/-log2FoldChange and an adjusted p-value <0.05. Each point represents a single gene and is colored were annotated by a combination of KEGG pathways and manual classification as spore-associated. For figure (B) each point represents the mean of all biological replicates for each treatment, and error bars represent SEM. Figures (C-F) show each bar representing the mean of all biological replicates for each treatment, and error bars represent the SEM. ** p < 0.01, *** p < 0.001.

We next sought to determine if the transcriptional response of the pathogen which could help explain the decrease in virulence in cocolonized mice, so we performed RNAseq on *C. difficile* from mouse cecal content collected at day two and used DESeq2 for differential expression analysis. We observed 95 significantly differentially expressed genes (89 down and 6 up) in *C. difficile* in cocolonized mice compared to monocolonized (Figure 4H, log2 fold change ± 2 and adjusted p-value < 0.05). Manual annotation of gene functions showed that 50/89 significantly decreased genes were involved in germination and sporulation pathways (Table S1) ^45,46^. Specifically, *sinR* and *sinR’,* two regulators of key virulence pathways affecting toxin, motility, and sporulation, were decreased in *C. difficile* (Table S1)^47^. This suggests that the effects of *C. hiranonis* on *C. difficile in vivo* may be in part due to transcriptional effects on genes essential for toxin and spore activity, which are essential to virulence.

### *C. hiranonis* produces a subinhibitory concentration of DCA in the presence of CA to suppress *C. difficile* virulence *in vitro* and *in vivo* without affecting colonization

*C. hiranonis* can decrease *C. difficile* toxin activity and spores *in vivo*, but the mechanism by which this occurs is still unclear. We hypothesized that BAs made in the presence of CA could be responsible for the suppression of *C. difficile* toxin activity and spores. To test this hypothesis, we again leveraged targeted BA metabolomics to define the BAs present in mice cocolonized with *C. hiranonis* and *C. difficile* compared to those monocolonized with *C. difficile,* as well as all other treatments. Principal component analysis (PCA) showed that *C. hiranonis* and *C. difficile* cocolonized mice at days 2 and 4 post *C. difficile* challenge clustered distinctly from all other samples (Figure 5A). CA, DCA, and 7-oxoDCA were shared between these mice and *in vitro* samples from *C. difficile* grown in *C. hiranonis* and CA spent media, which all had suppressed toxin activity (Figure 5B; Figure S3A-C). Together, this indicates that *C. hiranonis* modifies CA into different BAs, and those that overlap *in vitro* and *in vivo* may be affecting *C. difficile* virulence.

**Figure 5.**
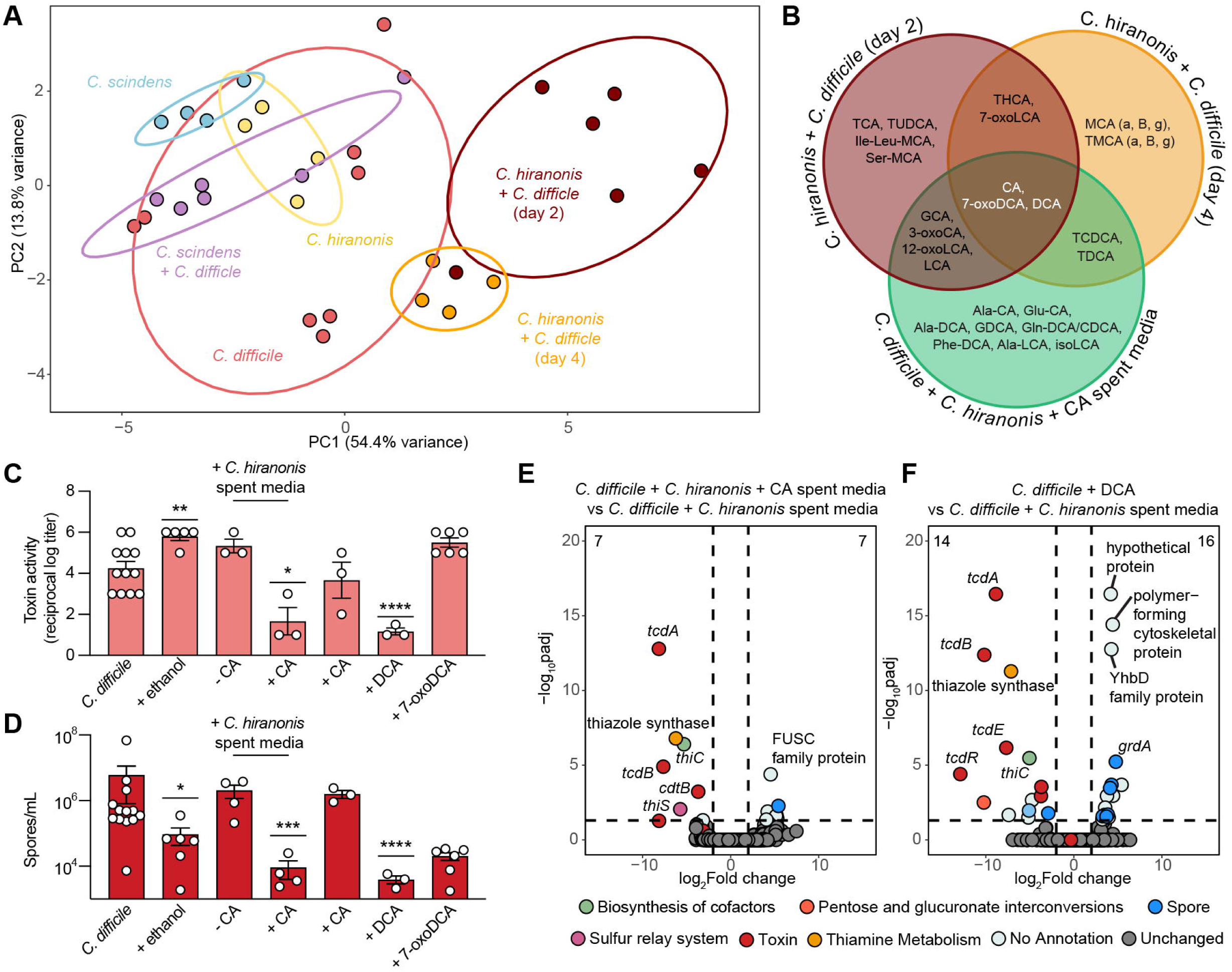
Subinhibitory concentrations of DCA reduce *C. difficile* virulence through decreased toxin gene expression, toxin activity and spore formation *in vitro*. A) Principal component analysis of BA profiles of mice from Fig. 2, Fig. 4., and Figure S2F-H. (B) Venn diagram highlighting BAs significantly enriched *in vivo* in *C. hiranonis* + *C. difficile* mice at day 2 post challenge (red), day 4 post challenge (orange), and *in vitro* in *C. difficile* + *C. hiranonis* + 2.5 mM CA spent media (green). Significance in each condition was determined by t-test (log2 fold change > 2 and adjusted p value < 0.05). C) Toxin activity in *C. difficile* cultures with and without BAs after 24 hr of incubation in different conditions. Statistical significance was determined by performing a Kruskal-Wallis test, comparing each condition to the pure culture of *C. difficile*. D) Spore load of *C. difficile* cultures with and without BAs after 72 hr of incubation in different conditions. Statistical significance was determined by performing a Kruskal-Wallis test on transformed data, comparing each condition to the pure culture of *C. difficile*. E) Volcano plots showing change in *C. difficile* gene expression *in vitro* in cultures grown in *C. hiranonis* and 2.5 mM CA compared to *C. difficile* and *C. hiranonis* spent media or (F) *C. difficile* and 0.625 mM DCA compared to *C. difficile* and *C. hiranonis* spent media. Differential gene expression was determined by DESeq2, and significance was defined as +/- log2FoldChange of 2 and an adjusted p- value <0.05. Genes were annotated by a combination of KEGG pathways and manual classification as spore-associated. * p < 0.05, ** p < 0.01, *** p < 0.001, **** p < 0.0001.

Since CA, DCA, and 7-oxoDCA are present in conditions where *C. difficile* virulence is decreased, we hypothesized that one or more of these BAs is responsible for this phenotype. We grew *C. difficile* in pure culture supplemented with each BA at a subinhibitory concentration of 0.625 mM. We also included a culture of *C. difficile* grown in *C. hiranonis* supplemented with CA spent media as a positive control, and saw that, similarly to Figure 3, the spent media decreases both toxin activity (Figure 5C Welch’s t-test, p = 0.039) and spores (Figure 5D, Welch’s t-test p = 0.0001) compared to *C. difficile* alone. In comparison, *C. hiranonis* spent media without CA does not affect toxin activity or spores (Figure 5C-D). 7-oxoDCA did not affect toxin activity when compared to its ethanol vehicle control, which increased toxin activity (Figure 5C, Welch’s t-test p = 0.0011) and decreased spore load (Figure 5D, Welch’s t-test p = 0.0132). The only BA that mirrored the effects of *C. hiranonis* and CA spent media was DCA (Figure 5C-D Welch’s t-test, p < 0.0001). We also tested a lower concentration of each BA, including DCA, (0.078 mM) and found that the effects on toxin activity and spores were less pronounced, suggesting that the concentration of DCA made is important for growth-independent suppression of virulence (Figure S3D-I).

Secondary BAs can also bind to *C. difficile* toxin, altering its conformation, which prevents binding to host cellular receptors and subsequent tissue damage ^48,49^. We next sought to determine how these BAs might be altering *C. difficile* toxin activity by first testing if they were able to directly bind to the toxin itself. We titrated DCA, 7-oxoDCA, and the control methyl-cholate (meCA) and used differential scanning fluorometry (DSF) to determine the binding EC_50_ of each BA. Binding occurred with an EC_50_ value of 179 μM for DCA and at 202 μM for 7-oxoDCA, while the positive control and strong binder meCA bound at 20 μM (Figure S3G). We also titrated each BA and measured cell rounding protection and found that DCA and 7-oxoDCA weakly protect against cell rounding when compared to meCA, which is also a robust rounding inhibitor (Figure S3H). Our binding EC_50_ and cell rounding protection assays indicate that the BAs produced by *C. hiranonis* in the presence of CA do not prevent *C. difficile* toxin mediated tissue damage by toxin binding alone.

As none of the BAs were able to bind to toxin, we next wanted to investigate whether they modulated *C. difficile*’s transcriptional response. We performed RNAseq on the *C. difficile* cultures from Figure 5C-D. We found that *C. difficile* grown in spent media from *C. hiranonis* and 0.625 mM CA had decreased expression of the toxin genes *tcdA, tcdB,* and *cdtB* relative to *C. difficile* grown in spent media from *C. hiranonis* without CA (Figure 5E, log2 fold change ± 2 and adjusted p-value < 0.05; Supplemental Table 2). Likewise, we observed a similar decrease in *tcdA, tcdB, tcdE,* and *tcdR* expression in *C. difficile* grown in DCA compared to *C. difficile* grown in *C. hiranonis* spent media (Figure 5F, log2 fold change ± 2 and adjusted p-value < 0.05; Table S2).

We also compared gene expression in *C. difficile* cultures in Figure 5C and 5D to *C. difficile* grown alone to determine whether these BAs were also able to affect *C. difficile* metabolism. *C. hiranonis* and CA spent media increased the expression of genes associated with many metabolic pathways including ABC transporters, butanoate metabolism, phosphotransferase systems, and starch and sucrose metabolism as well as spore-associated pathways, while amino acid metabolism genes are decreased (Figure S4B, Table S2). DCA causes a similar change in spore-associated genes and genes involved in phosphotransferase systems, while CA does not have an effect on *C. difficile* gene expression (Figure S4D, Table 2). Other BAs from Figure 5C and 5D had a minimal effect on *C. difficile*’s metabolism (Figure S4D-F, Table S2). This suggests that *C. hiranonis*-produced DCA at subinhibitory concentrations alters toxin expression, thereby decreasing both toxin activity and virulence.

### The addition of *C. hiranonis* BSH allows *C. scindens* to provide partial protection against toxin mediated damage from CDI

The presence of an encoded BSH in *C. hiranonis* and the presence of DCA in mice given *C. hiranonis*, but not in *C. scindens*, led us to hypothesize that BSH activity is essential for releasing the CA from TCA, which is a necessary step for secondary BA production *in vivo*. BSHs belong to the larger N-terminal nucleophile (Ntn)-hydrolase enzyme superfamily, where their primary function is deconjugation of glycine or taurine-conjugated primary BAs, releasing the unconjugated BA and free amino acid. Recently, our lab showed a conserved substrate selectivity loop containing a motif dictating a preference for glycine (Y-S-R-X) or taurine-conjugated (G-X-G) BAs ^30^. *C. hiranonis* BSH (ChirBSH) contains the glycine-preferring motif (Figure 6A), leading us to test the substrate specificity and activity of the enzyme *in vitro*. Following recombinant protein production and purification (Figure 6B), we isolated ChirBSH and measured the activity with the primary BAs GCA, GCDCA, TCA, and TCDCA and the secondary BAs GDCA and TDCA. ChirBSH exhibits a preference for glycine-conjugated BAs, particularly GDCA, but it still has appreciable activity with taurine-conjugates including TCA (Figure 6C).

**Figure 6.**
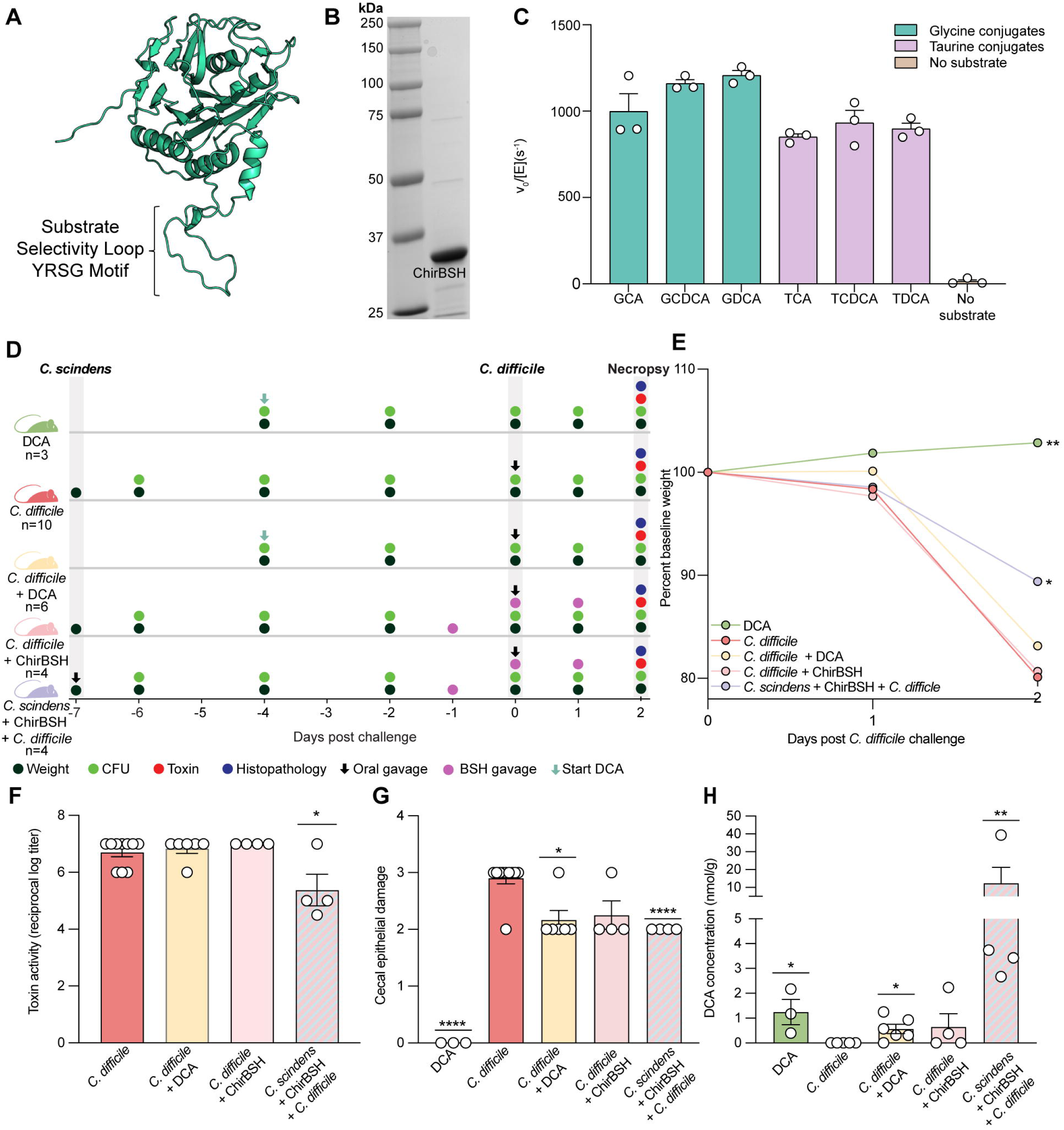
*C. hiranonis* BSH provides partial protection against disease in mice cocolonized with *C. scindens* and *C. difficile*. A) AlphaFold2 model of *C. hiranonis* BSH (ChirBSH) with the YSRG motif highlighted in the substrate specificity loop. B) 10% SDS-PAGE of ChirBSH following recombinant production and purification. C) Ninhydrin activity assay showing purified *C. hiranonis* BSH activity with glycine- or taurine-conjugated substrates *in vitro*. D) Experimental overview of gnotobiotic animal model. E) Percent change in weight from baseline was tracked from days 0-2 post *C. difficile* challenge in mice supplemented with DCA (n = 3), *C. difficile* monocolonized mice (n = 10), *C. difficile* monocolonized mice supplemented with DCA (n = 6), *C. difficile* monocolonized mice with ChirBSH (n = 4), and *C. difficile* and *C. scindens* cocolonized mice with ChirBSH (n = 4). Statistical significance was determined by using Kruskal-Wallis test with Dunn’s correction to compare all conditions to *C. difficile* monocolonized mice. F) *C. difficile* toxin activity measured by Vero cell cytotoxicity assay on cecal contents from (E). Statistical significance was determined by comparing *C. difficile* monocolonized mice to cocolonized mice with a Mann-Whitney test. G) Histopathological analysis of cecal epithelial damage at day 2. Tissues were scored on a scale from 0-4 and statistical comparisons by comparing the score of each condition to that of *C. difficile* monocolonized mice by two-way ANOVA. H) Concentration of DCA in cecal content of mice at day 2. Statistical significance was determined by Mann-Whitney test, comparing the concentration of DCA to that of *C. difficile* monocolonized mice. * p < 0.05, ** p < 0.01, **** p < 0.0001.

To test if the BSH is needed to provide a protective effect with *C. scindens in vivo*, mice were stably colonized with *C. scindens* for seven days before 10 μg of purified ChirBSH recombinant enzyme was administered via oral gavage on day -1 before challenge with *C. difficile* spores on day 0 (Figure 6D, Figure SA-B). Mice cocolonized with *C. scindens* and *C. difficile* and given ChirBSH lost less weight than mice monocolonized with *C. difficile* (Figure 6E, Kruskal-Wallis test p = 0.0.0111). Additionally, *C. difficile* bacterial load was higher in monocolonized mice that received ChirBSH (Figure S5C, Mann-Whitney test, p = 0.0360). While ChirBSH challenge did not affect *C. difficile* toxin activity in monocolonized mice, cocolonized mice with ChirBSH had less toxin activity (Figure 6F, Mann-Whitney test p = 0.0210) and epithelial damage than *C. difficile* monocolonized mice (Figure 6G, Two-way ANOVA with Dunnett’s multiple comparison correction p <0.0001). Cecal edema and inflammation did not change across treatments, and colonic histopathology was consistent with mice monocolonized with *C. difficile* (Figure S5D-E). Cocolonized mice given ChirBSH had higher DCA in their cecum compared to the other groups (Figure 6H, Mann-Whitney test p = 0.0079). While DCA was increased in cocolonized mice given ChirBSH, there were minimal effects on the rest of the BA pool (Figure S5F). These findings indicate that giving ChirBSH to mice cocolonized with *C. scindens* and *C. difficile* can provide partial protection against toxin mediated damage from CDI and allow for the liberation of CA to make DCA.

We also wanted to test whether supplementing excess DCA alone would elicit a protective effect similar to that observed in mice cocolonized with *C. scindens* and *C. difficile* that were given ChirBSH. To do so, we supplemented the drinking water of GF mice with 1.25 mM DCA starting on day -4 before challenging a subset of mice with *C. difficile* spores on day 0 (Figure 6D, Figure S5A-B). Supplementing DCA in drinking water did not prevent weight loss, colonization or toxin activity after *C. difficile* challenge (Figure 6E-F, Figure S5C). However, supplementing DCA to *C. difficile* colonized mice did decrease cecal epithelial damage, but not inflammation and edema or colonic tissue damage (Figure 6G, two-way ANOVA p = 0.0.0167, Figure S5D-E). DCA supplementation also increased the amount of DCA detected in the cecum and to some extent CDCA (Figure 6H, Mann-Whitney test p = 0.0179, Figure S5G) but did not alter other BAs like TDCA or GDCA (Figure S5F). The amount of DCA in these mice was determined to be less than that of *C. scindens* and *C. difficile* cocolonized mice given ChirBSH (Figure 6H, Mann-Whitney test p = 0.0095). While exogenous DCA supplementation did cause a decrease in epithelial damage, it did not affect *C. difficile* toxin activity or significantly increase the amount of free DCA in a manner like that of *C. scindens* and *C. difficile* cocolonized mice given ChirBSH.

## DISCUSSION

To disentangle the relative contributions of BA modifications versus nutrient competition to Clostridia-mediated colonization resistance (CR) against *C. difficile*, we conducted a series of complementary *in vitro* and *in vivo* experiments. We found that neither *C. scindens* nor *C. hiranonis* altered *C. difficile* colonization or virulence through competition for nutrients alone. However, when supplemented with the primary bile acid CA, *C. hiranonis* generated subinhibitory concentrations of the secondary bile acid DCA that attenuated *C. difficile* virulence without preventing colonization. In contrast, CA-supplemented *C. scindens* generated inhibitory concentrations of DCA *in vitro* that reduced *C. difficile* growth and hence toxin activity but failed to produce DCA *in vivo* and consequently did not protect against colonization or virulence. Neither organism was able to alter *C. difficile* growth or virulence on their own in the absence of CA, indicating that competition for nutrients alone with these Clostridia strains does not provide CR. Using a bottom-up approach, we found that commensal-mediated effects on *C. difficile* are dependent on strain, starting BA substrate concentration, and context.

Metabolic context is especially important when comparing the behavior of *C. scindens* and *C. hiranonis* in the presence of BAs as the effect on *C. difficile* varied by organism and CA concentration. For *C. scindens*, low dose CA only reduced toxin activity, whereas a high dose reduced both growth and toxin activity (Figure 1B, 1D). In contrast, *C. hiranonis* consistently reduced *C. difficile* toxin activity, but not growth, regardless of CA concentration (Figure 3B, 3D). These divergent patterns likely reflect differences in microbial responses to CA, tolerance to DCA, or regulation of the *bai* operon. Although both organisms increase *bai* expression when 2.5 mM CA is added, expression in *C. scindens bai* is roughly ∼10x higher than that of *C. hiranonis* ^41^. This is consistent with *C. scindens’* more efficient conversion of CA to DCA (1.97 mM from 2.5 mM CA, versus 0.78 mM for *C. hiranonis*) and greater DCA tolerance (MIC 1.88 mM, versus 0.78 mM for *C. hiranonis*) ^41^. This may explain why *C. hiranonis* produces subinhibitory concentrations of DCA, as it has a lower MIC for DCA. These low concentrations of DCA are sufficient to reduce toxin activity but are insufficient to inhibit *C. difficile* growth and potentially represent a mechanism of self-preservation by *C. hiranonis*. Alternatively, because *C. hiranonis* encodes a BSH, it may reconjugate DCA to amino acids, reducing DCA levels that are toxic to itself, as was observed *in vitro* with alanine-, glycine-, glutamine-, and phenylalanine-conjugated DCA^38^ (Figure 5B, Supplemental Table 2). This could represent an elegant and underappreciated protective mechanism that bacteria can utilize in the presence of toxic BA levels, as the ability of BAs to conjugate with non-canonical amino acids is a relatively new finding^30,50,51^. The presence of an encoded BSH in *C. hiranonis* also enables access to host-conjugated TCA for 7α-dehydroxylation *in vivo*; in comparison *C. scindens*, which lacks a BSH, did not produce DCA *in vivo* (Figure S1C). Together, these observations suggest that *bai* operon activity, BA tolerance, and accessory BA-modifying functions jointly dictate the magnitude and form of secondary BA output. In this study, the BSH encoded by *C. hiranonis* was essential for secondary BA production or DCA *in vivo*.

It has been well established that *C. difficile* is sensitive to changes in the BA pool. Primary BAs like TCA are strong germinants of *C. difficile* spores and are increased after antibiotic use and during CDI ^4,52^. In comparison, secondary BAs are associated with restoration of CR and reduce *C. difficile* spore germination, growth, toxin expression, and toxin activity ^53–56^. DCA can induce biofilm formation and act as a germinant, although it blocks vegetative cell outgrowth, indicating that there is nuance in the effects of secondary BAs on the different stages of the *C. difficile* lifecycle ^52,57^. Secondary BAs are also able to bind *C. difficile* toxin, with LCA being an especially strong binder and DCA binding with less affinity ^48^. DCA also directly alters the transcription of *tcdA* and *tcdB*, which may act as a second mechanism of protection against *C. difficile* toxin^57^. In this study we show that subinhibitory concentrations of DCA decrease virulence factors including toxin expression, activity and spore formation. We also know that many BAs can directly affect the host response. DCA is an important regulator of host BA metabolism, and does so by modulating synthesis via nuclear farnesoid X receptor (FXR), while UDCA has been shown to also limit inflammation during CDI by altering immune receptors including G-protein-coupled membrane receptor 5 (TGR5) ^58,59^. Although DCA supplementation decreased epithelial damage, it did not reach levels *in vivo* that directly affected *C. difficile* toxin activity as observed in *C. hiranonis* cocolonized mice or *C. difficile* and *C. scindens* cocolonized mice given ChirBSH. These findings suggest that delivery of subinhibitory concentrations of DCA to the large intestine might be better coming from bacteria and enzymes.

While secondary BAs have long been considered drivers of CR, other BA classes play equally important roles in maintaining protection against CDI. Epimers of LCA such as isoLCA and isoalloLCA are produced by HSDH and have also been shown to decrease *C. difficile* growth and toxin expression in addition to influencing host immune cell development ^55,60,61^. In comparison, 7-oxoDCA is also made via HSDH activity but was not found to impact *C. difficile* growth, spore load, or toxin activity (Figure 5B-C, Figure S3). Importantly, 7-oxoDCA is made by *C. difficile* through its 7 ⍰ HSDH, indicating that *C. difficile* makes its own BA, although it is not yet known whether this benefits *C. difficile* ^43^. Some strains of *C. difficile in vitro* have been shown to display BSH-like activity, although there is no annotated *bsh* in the genome^62^. We did not see this in our study as we used *C. difficile* R20291. Newly identified MCBAs have also been found to influence *C. difficile* growth and virulence and increase the overall complexity of the BA pool ^30^. As our understanding of the BA pool advances^63,64^, it will be important to also consider how BAs impact the *C. difficile* lifecycle, the microbiota and the host.

Although BA metabolism emerged as the dominant driver of virulence attenuation in our experiments, it nonetheless intersected with *C. difficile* metabolism. Nutrient availability is important for CR especially in more complex models ^23,33–35,65–67^. *C. scindens* and *C. hiranonis* both utilize Stickland fermentation and consume amino acids like glycine, proline, serine, and threonine, which *C. difficile* also uses ^37,68^. Since the specific amino acid needs of *C. scindens* and *C. hiranonis* overlap with those of *C. difficile*, amino acids are a point of competition between these organisms and may be a contributing factor to CR ^36,38^. While *C. scindens* has been shown to compete with *C. difficile* for proline *in vitro*, carbohydrate utilization pathways in *C. difficile* are also differentially expressed during coculture ^36^. Similar studies have shown that *C. difficile* can utilize alternative sugar alcohol metabolic pathways during competition with *P. bifermentans*, suggesting that carbohydrate utilization may also be important ^33^. This aligns with our work, which shows *C. hiranonis* and CA spent media cause changes in expression of genes associated with *C. difficile* carbohydrate metabolism (Figure S4B). Although we did not see complete inhibition of *C. difficile* as in other studies, we did observe a decrease in *C. difficile* virulence in the presence of *C. hiranonis* and CA spent media *in vitro* and *C. hiranonis in vivo*, suggesting that products from *C. hiranonis* may be driving metabolic shifts in *C. difficile* that translate to reduced virulence. Moreover, prior work has linked amino acid and carbohydrate availability to BA metabolism ^69,70^, hinting at a more complex interplay between nutrient landscapes and microbial BA transformations in mediating disease prevention. While we have considered competition for nutrients in the context of preventing *C. difficile* colonization and virulence, others have shown that, in certain instances, cross-feeding enhances virulence and worsens disease outcomes. Specifically, organisms such as Enterococci increase *C. difficile* virulence by a combination of cross-feeding ornithine, which can be converted to proline, while simultaneously restricting arginine availability^71^. This further increases the complexity, and highlights how understanding the role and importance of both BA metabolism, nutrient competition and cross-feeding in more complex models will be important as we continue to define the different mechanisms of CR against *C. difficile*.

These mechanistic insights have direct implications for therapeutic design. We demonstrate that protection against virulence *in vivo* can be conferred by Clostridia encoding specific sets of BA-altering genes, specifically BSH and the *bai* operon, and that direct supplementation with secondary BAs such as DCA provides partial protection against toxin-mediated tissue damage (Figure 5K). While this route may seem promising, DCA supplementation did not prevent clinical signs of disease or decrease toxin activity (Figure 4B-D and 5H-J), potentially due to dosing or host conjugation to TDCA, which may limit bioavailability. One potential solution is modifying BAs to restrict host absorption while preserving luminal activity, a recent approach that has shown promise *in vivo*, but is still in early stages of research and currently restricted to LCA ^49^. In parallel with small molecules, our findings also motivate organism and enzyme-based therapeutic approaches. We show that supplementing purified ChirBSH provides partial protection against disease *in vivo* when paired with a 7[7]-dehydroxylating bacterium (Figure 6), consistent with prior work demonstrating that BSH cocktails from lactobacilli partially inhibit *C. difficile* by producing MCBAs ^30^. While enzyme-based approaches will require optimization of dosing and delivery, our data suggests that *C. hiranonis* itself represents a particularly compelling candidate organism for a defined consortium. Although it does not block colonization, *C. hiranonis* encodes both BSH and *bai* operon genes, enabling production of a broad spectrum of secondary BA metabolites, including those implicated in the efficacy of Vowst and Rebyota, therapeutics that are currently being used to treat rCDI patients ^14,15,38^. Moreover, *C. hiranonis* has been included in a synthetic community associated with inhibiting *C. difficile* growth *in vitro* and been associated with prevention of *C. difficile* carriage in companion animals ^44,68^. Thus, its ability to produce subinhibitory amounts of DCA that attenuates toxin activity without suppressing growth, and limiting host tissue damage, make it especially well-suited for virulence-targeted microbiome therapeutics.

Critically, our findings diverge from recent work, which conclude that nutrient competition with organisms such as *C. scindens* or *C. hiranonis*, as opposed to bile acid modulation, confers a protective effect against CDI in a gnotobiotic mouse model^23,34,36^. However, it is important to distinguish between the types of disease metrics used in those studies as opposed to our own. Specifically, previous studies rely upon weight loss and percent survival as a primary measurement of CR, whereas we approach this with a more nuanced range of metrics including measuring weight loss, bacterial enumeration by plating on BHI medium, toxin activity, spores, and disease by measuring histopathological changes to the gut tissue. By incorporating these measures, we show that while weight is important, other metrics are more informative. In addition, some previous studies use a strain of *C. scindens* that has an antimicrobial peptide, *C. scindens* ATCC 35704, which may cause confounding effects that bias the results in favor of competition of nutrients. In contrast, our work suggests that while competition for nutrients may have some effect on *C. difficile,* as evidenced by changes in *C. difficile* gene expression in the presence of *C. hiranonis* and CA spent media, the effects of BA modulation have far more significant effects on disease metrics in this model. Interestingly, our finding that subinhibitory concentrations of secondary BAs can decrease toxin-mediated disease independently of colonization further calls into question the extent to which competition for nutrients influences CR in this model. As such, we urge the field to reconsider the dogma of nutrient competition as the primary or sole factor contributing to CR and examine existing data with more nuance and care for the specific microbial strains, mouse models, and disease outcomes measured.

Despite the rigorous and clear mechanisms proposed, as evidenced by our paired *in vitro* and *in vivo* approaches, several limitations warrant consideration. First, due to the limited genetic toolkits available for Clostridia, we were unable to truly separate BA metabolism from amino acid metabolism and thus investigate these aspects separately. Additionally, we only considered two strains of commensals, *C. scindens* VPI12708 and *C. hiranonis* TO-931, and one *C. difficile* strain, R20291, therefore other strains may exhibit distinct metabolic capabilities or interaction dynamics. For example, *C. scindens* ATCC 35704 produces a tryptophan-derived antibiotic that can inhibit *C. difficile*, altering growth on its own ^72^, and *C. hylemonae* has been proposed to preferentially utilize CDCA instead of CA^41^. Likewise, different *C. difficile* strains may vary in their sensitivity to BA-mediated modulation. Methodological constraints may also have shaped the scope of our conclusions, as we used a single rich medium (BHI) to support robust growth of all organisms *in vitro*, as opposed to a defined medium. Furthermore, we only supplemented CA in the context of the *bai* and not did not test the effects of CDCA and its product LCA. Moreover, while our gnotobiotic mouse model is powerful tool to test microbial mechanistic dynamics *in vivo*, it does not fully approximate host-microbiota complexity. Finally, we tested only single dosages of DCA and BSH enzyme supplementation, and future work will be necessary to determine the optimal dosing strategies of each.

Looking forward, extending these findings to more complex ecological contexts will be essential. Future studies should examine how *C. scindens* and *C. hiranonis* interact with *C. difficile* in a consortium, an antibiotic treated mouse model and within a defined human consortium such as hCom1 ^73^. Understanding how these microbes’ function within communities and how their metabolic outputs scale across microbial ecosystems will be critical for translating mechanistic insights into therapeutic strategies. Further, repeating the *in vitro* studies from this work using defined amino acid or carbohydrate media will help clarify how nutrient availability shapes BA metabolism and vice versa.

Collectively, these findings challenge the prevailing assumption that CR should be the solitary objective of microbiome-based therapeutics and instead suggest that virulence attenuation may represent an additional clinically meaningful outcome. They also underscore how host metabolic context profoundly shapes microbial function, such that the same microbe can exert fundamentally different effects on disease outcomes depending on substrate availability. More broadly, these findings also underscore the need to consider microbial metabolic activity and not just community composition or pathogen load when defining mechanisms of CR and designing microbiome-based therapeutics. In this framework, *C. hiranonis* emerges as promising member of a rationally designed consortia when nutritional, host, and metabolic contexts are carefully considered.

### Limitations of the study

This is included in the discussion section.

## RESOURCE AVAILABILITY

### Lead contact

Further information and requests for resources and reagents should be directed to and will be fulfilled by the lead contact, Casey M. Theriot.

### Materials availability

Raw sequences from RNA sequencing data have been deposited in the Sequence Read Archive (SRA) under BioProject ID: PRJNA1435196. Data acquired from targeted metabolomics has been deposited in MASSive under MSV000100897. Bile Acid LC-IMS-MS library (negative mode) from Zenodo- Baker lab can be found here: https://doi.org/10.5281/zenodo.15318635. Any additional information required to reanalyze the data reported in this paper is available from the lead contact upon request.

## ACKNOWLEDGMENTS

C.M.T. is funded by the National Institute of General Medical Sciences of the National Institutes of Health under award number R35GM119438 and R35GM149222. The Gnotobiotic Core at the College of Veterinary Medicine, North Carolina State University is supported by the National Institutes of Health funded Center for Gastrointestinal Biology and Disease, NIH P30 DK034987. S.C.K. is funded by the National Institutes of Health under award number 1T32GM133366, the NCSU Genetics and Genomics Scholars Program, and the L.W. Parks Endowment Microbiology Research Award. E.S.B. is funded by research grants from the National Institutes of Health (P42 ES027704, R01 GM141277, and RM1 GM145416), and the National Science Foundation. The views expressed in this manuscript do not reflect those of the funding agencies.

We thank Jason Ridlon for providing the commensal Clostridia strains and Trevor Lawley for providing the *C. difficile* strain used in these experiments. The authors also acknowledge the Roy J. Carver Biotechnology Center at the University of Illinois at Urbana-Champaign for providing RNA sequencing and the McGuire VA Medical Center LC-MS Core Lab at Virginia Commonwealth University for providing LC-MS analysis.

## AUTHOR CONTRIBUTIONS

Conceptualization, S.C.K and C.M.T.; methodology, S.C.K, C.E.P., A.S.G., N.S.L., S.A.T., E.C.V., G.Z., J.T., R.M., E.C.R., E.S.B., C.M.T.; Investigation, S.C.K, C.E.P., A.S.G., N.S.L., S.A.T., E.C.V., G.Z., J.T., E.C.R.; writing—original draft, S.C.K. and C.M.T.; writing—review & editing, S.C.K., A.S.G., and C.M.T; funding acquisition, S.C.K. and C.M.T.; resources, C.M.T.; supervision, E.S.B., R.M, and C.M.T.

## DECLARATION OF GENERATIVE AI AND AI-ASSISTED TECHNOLOGIES

Nothing to declare.

## SUPPLEMENTAL INFORMATION

**Document S1. Figures S1–S5 and Table S1-S2**

**Figure S1. *C. scindens* does not prevent *C. difficile* colonization and disease *in vivo* even when cholate is supplemented in drinking water**

A) Percent change in weight from baseline was tracked from days -7 to 0 post *C. difficile* challenge. B) Bacterial load in fecal content collected from days -7 to 0 post *C. difficile* challenge. C) Histopathological changes to colonic tissue from each treatment. Tissues were scored for edema, epithelial damage, and inflammation on a scale of 0-4 for a maximum total score of 12 and statistical comparisons were made between each scoring category across all conditions by two-way ANOVA (a, p < 0.015 edema; b, p < 0.001 epithelial damage; c, p < 0.0001 epithelial damage; d, p < 0.0001 inflammation). D) Representative photomicrographs of colon tissue collected from mice in (C). E) Volcano plot showing change in BA enrichment in *C. scindens* + *C. difficile* cocolonized mice compared to *C. difficile* monocolonized mice. Significance was determined by t-test. F) Volcano plot showing change in BA enrichment in *C. difficile* monocolonized mice compared to GF mice. Significance was determined by t-test. G). Percent change in weight from baseline was tracked from days 0-2 post *C. difficile* challenge in *C. scindens* monocolonized mice + CA (n = 3), *C. difficile* monocolonized mice + CA (n = 4), and cocolonized mice + CA (n = 3). Statistical significance was determined by using Kruskal-Wallis test with Dunn’s correction to compare each condition to the *C. difficile* monocolonized mice (**, p < 0.01). H) *C. difficile* bacterial load in cecal content collected at day 2. I) *C. difficile* toxin activity measured by Vero cell cytotoxicity assay on cecal content from mice in (G).

**Figure S2. *C. hiranonis* ameliorates disease *in vivo* with and without cholate supplemented in drinking water and up until day 4 post *C. difficile* challenge**

A) Percent change in weight from baseline was tracked from days -7 to 0 post *C. difficile* challenge. B) Bacterial load in fecal content collected from days -7 to 0 post *C. difficile* challenge. C) Histopathological changes to colonic tissue from each treatment. Tissues were scored for edema, epithelial damage, and inflammation on a scale of 0-4 for a maximum total score of 12 and statistical comparisons were made between each scoring category across all conditions by two-way ANOVA (a, p < 0.05 edema; b, p < 0.001 epithelial damage; c, p < 0.001 epithelial damage; d, p < 0.0001 epithelial damage; e, p < 0.0001 inflammation). D) Representative photomicrographs of colonic tissue collected from mice in (C). E) Percent change in weight from baseline was tracked from days 0-2 post *C. difficile* challenge in *C. hiranonis* monocolonized mice + CA (n = 3), *C. difficile* monocolonized mice + CA (n = 4), and cocolonized mice + CA (n = 3). Statistical significance was determined by using Kruskal-Wallis test with Dunn’s correction to compare each condition to the *C. difficile* monocolonized mice (**, p < 0.01). F) *C. difficile* bacterial load in cecal content collected at day 2. G) *C. difficile* toxin activity measured by Vero cell cytotoxicity assay on cecal content from mice in (E). Statistical significance was determined by Mann-Whitney test (*, p < 0.05). H) Percent change in weight from baseline was tracked from days 0-4 post *C. difficile* challenge in *C. difficile* monocolonized mice (n = 4), and *C. hiranonis* and *C. difficile* cocolonized mice (n = 4). Statistical significance was determined by using a Mann-Whitney test to compare each condition to the *C. difficile* monocolonized mice at days 2, 3, and 4 post challenge (*, p < 0.05). I) *C. difficile* bacterial load in cecal content collected at day 4 post challenge. J) *C. difficile* toxin activity measured by Vero cell cytotoxicity assay on cecal content from monocolonized mice collected at day 2 post challenge and cocolonized mice collected at day 4 post challenge. Statistical significance was determined by Mann-Whitney test (*, p < 0.05). K) *C. difficile* spore load from monocolonized (n = 6) and cocolonized mice (n = 4) collected at days 2 and 4 post challenge, respectively. Statistical significance was determined by Mann-Whitney test (*, p < 0.05). L) Histopathological changes to cecal tissue from each treatment. Tissues were scored for edema, epithelial damage, and inflammation on a scale of 0-4 for a maximum total score of 12 and statistical comparisons were made between each scoring category across all conditions by two-way ANOVA (a, p < 0.05 inflammation: b, p < 0.05 epithelial damage

**Figure S3. Bile acids shared across toxin attenuated phenotypes decrease toxin activity and spores *in vitro* but fail to bind to toxin B**

A-C) Volcano plots comparing BA pools across treatments, points are colored by significance and whether the BA is shared across A-C With A) comparing *C. hiranonis* and *C. difficile* cocolonized mice (day 2) to *C. difficile* monocolonized mice. (day 2), B) comparing *in vitro C. difficile* + *C. hiranonis* + CA spent media to *C. difficile* monoculture, and C) comparing comparing *C. hiranonis* and *C. difficile* cocolonized mice (day 4) to *C. difficile* monocolonized mice. (day 2). Vegetative (pink bars) growth and spore load (red bars) of *C. difficile* cultures at 24 hr (D), 48 hr (F), and 72 hr (H) of incubation. Corresponding toxin activity assay results from 24 hr (E), 48 hr (G), and 72 hr (I) of incubation. Each BA was tested at a low concentration (0.078 mM) and a high concentration (0.625 mM). J) Measurement of temperature-dependent fluorescence to calculate TcdB melting points in the presence of increasing concentrations of BAs. Comparison of 7-oxoDCA (pink circles), DCA (red circles) and methylcholate (blue circles) binding to TcdB by differential scanning fluorometry (DSF). Bars represent SEM of three independent experiments. K) Dose-dependent protection of TcdB-challenged IMR-90 cells by bile acids 7-oxoDCA (pink circles), DCA (red circles), and methylcholate (blue circles), with cell rounding measured by arrays can high-content imaging. Bars represent SEM of four independent experiments.

**Figure S4. *C. difficile* transcriptional response in the presence of different bile acids**

A-F) Volcano plots of DESeq2 output from *in vitro C. difficile* cultures. A) *C. difficile* + *C. hiranonis* spent media, B) *C. difficile* + *C. hiranonis* + CA spent media, C) *C. diffiicle* + 0.625 mM DCA, D) *C. difficile* + 0.625 mM CA, E) *C. difficile* + ethanol, or F) *C. difficile* + 0.625 mM 7-oxoDCA compared to *C. difficile* monoculture. Genes were annotated by KEGG pathway, and spores were annotated manually based on previous literature, as described in methods.

**Figure S5. Extended data from *C. hiranonis* BSH given to mice cocolonized with *C. scindens* and C. difficile**

A) Percent change in weight from baseline was tracked from days -7 to 0 post *C. difficile* challenge. B) *C. scindens* load in cecal content collected from days -7 to 0 post *C. difficile* challenge. C) Bacterial load in cecal content collected at day 2 post *C. difficile* challenge. Statistical significance was determined by comparing load to that of *C. difficile* monocolonized mice by Mann-Whitney test (*, p < 0.05; **, p < 0.01). D) Histopathological changes to cecal tissue of mice given DCA, *C. difficile* monocolonized mice, *C. difficile* monocolonized mice given DCA, *C. difficile* monocolonized mice given ChirBSH, or *C. scindens* and *C. difficile* cocolonized mice given ChirBSH. Tissues were scored for edema, epithelial damage, and inflammation on a scale of 0-4 for a maximum total score of 12 and statistical comparisons were made by comparing scores from each scoring category to the respective *C. difficile* monocolonized mouse score by two-way ANOVA ( a, p < 0.05 edema; b, p < 0.05 epithelial damage; c, p < 0.0001 epithelial damage; d, p < 0.05 inflammation). E) Histopathological changes to colonic tissue for each treatment. Tissues were scored for edema, epithelial damage, and inflammation on a scale of 0-4 for a maximum total score of 12 and statistical comparisons were made between each scoring category across all conditions by two-way ANOVA (a, p < 0.001 edema; b, p < 0.001 epithelial damage; c, p < 0.0001). F) Concentration of BAs in cecal content of mice given DCA (orange bars), *C. difficile* monocolonized mice given DCA (yellow bars), *C. difficile* monocolonized mice (pink bars), monocolonized mice given ChirBSH (light pink bars), *C. scindens* and *C. difficile* cocolonized mice (pink and blue striped bars), or cocolonized mice given ChirBSH (light pink and light blue striped bars). Statistical significance was determined by comparing bile acid concentrations to those of *C. difficile* monocolonized mice by a Mann-Whitney test (*, p < 0.05).

**Table S1:** *In vivo* RNAseq data in Figure 4H

**Table S2:** *In vitro* RNAseq data in Figures 5E-F, and Figure S4

## STAR⍰METHODS

### KEY RESOURCES TABLE

The items in the key resources table (KRT) must also be reported alongside the description of their use in the method details section. Literature cited within the KRT must be included in the references list. Please **do not edit the headings or add custom headings or subheadings** to the KRT. We highly recommend using <u>RRIDs</u> as the identifier for antibodies and model organisms in the KRT. To create the KRT, please use the template below or the <u>KRT webform</u>. See the more detailed <u>Word table template</u> document for examples of how to list items.

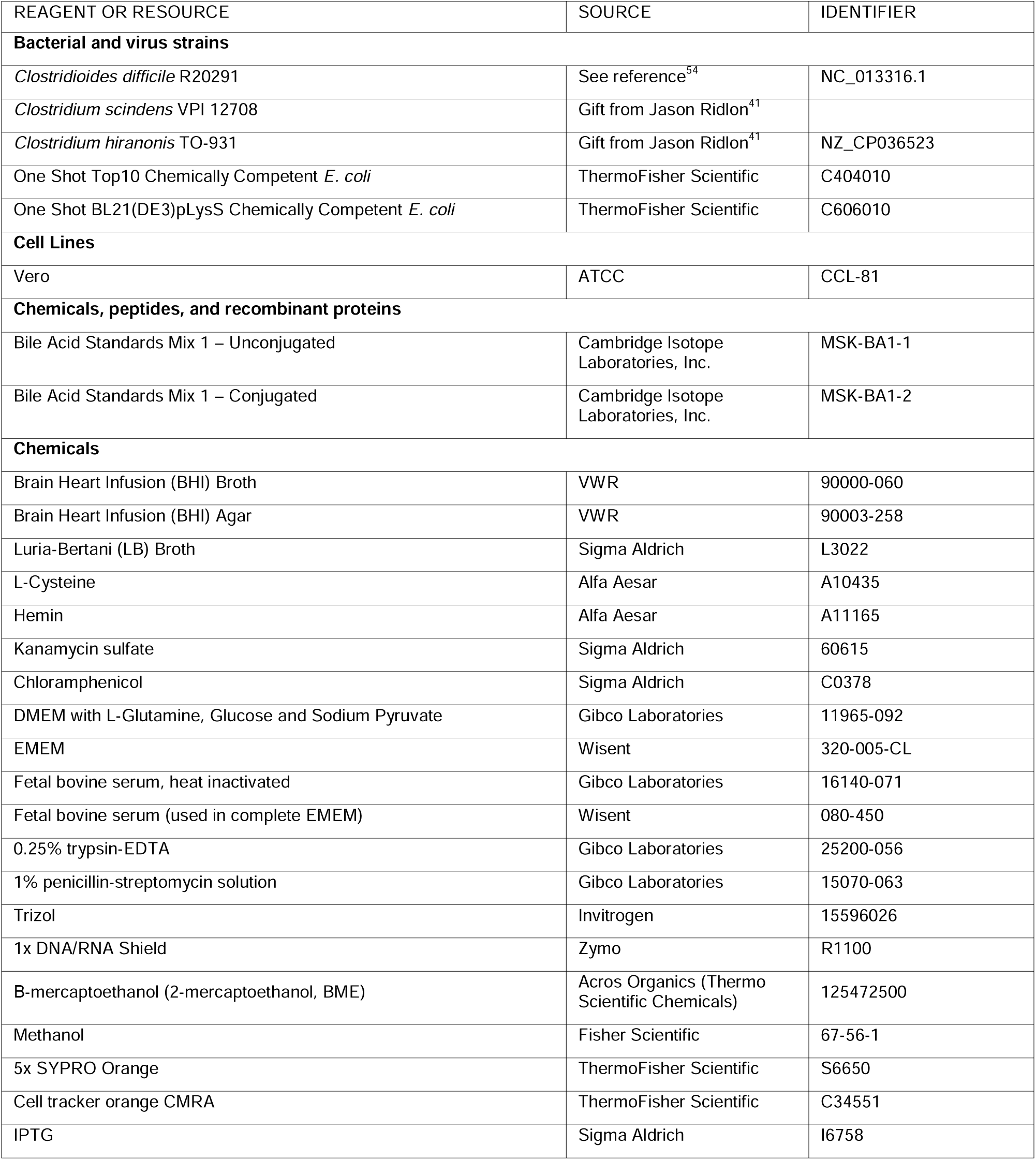
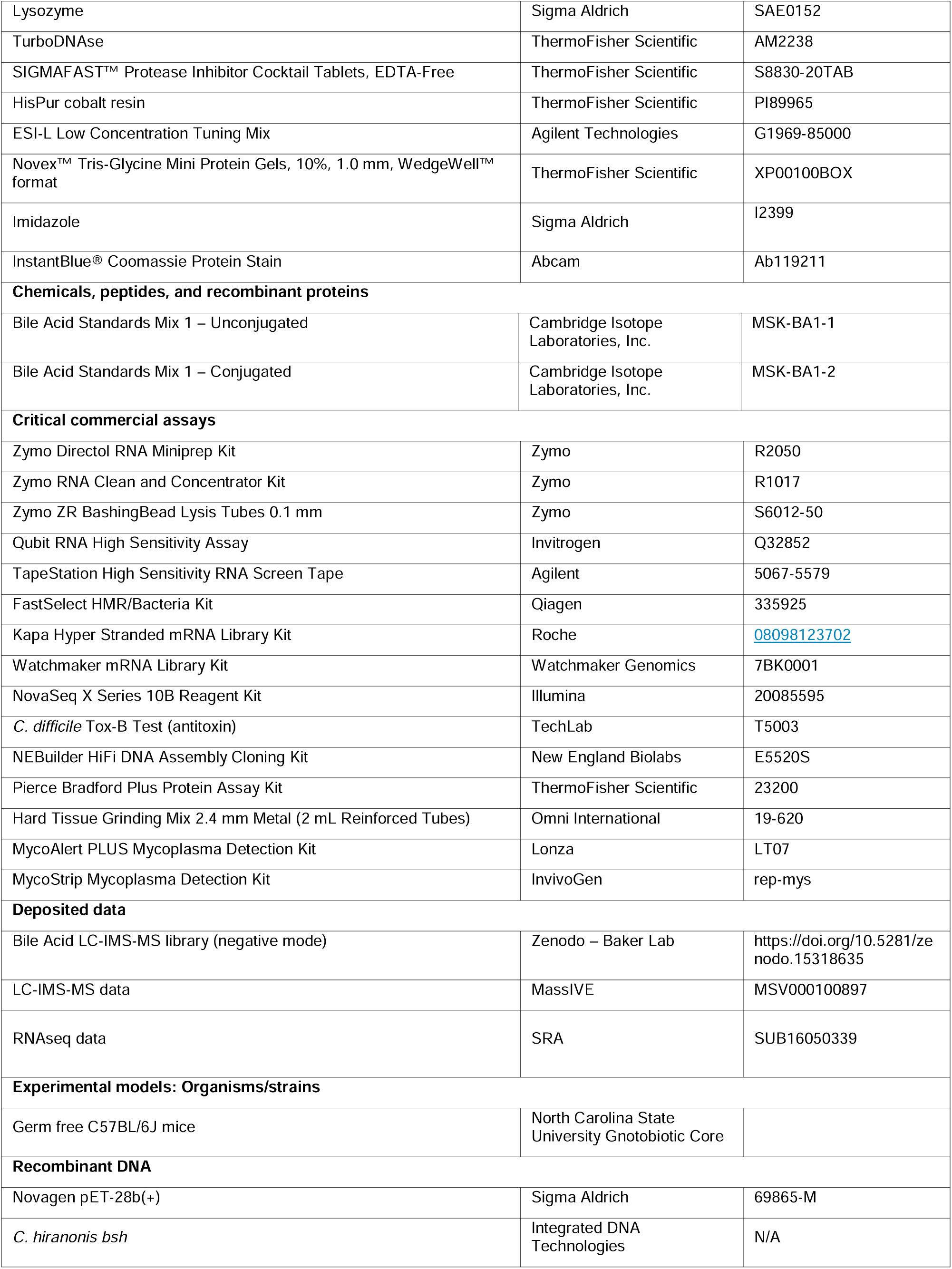
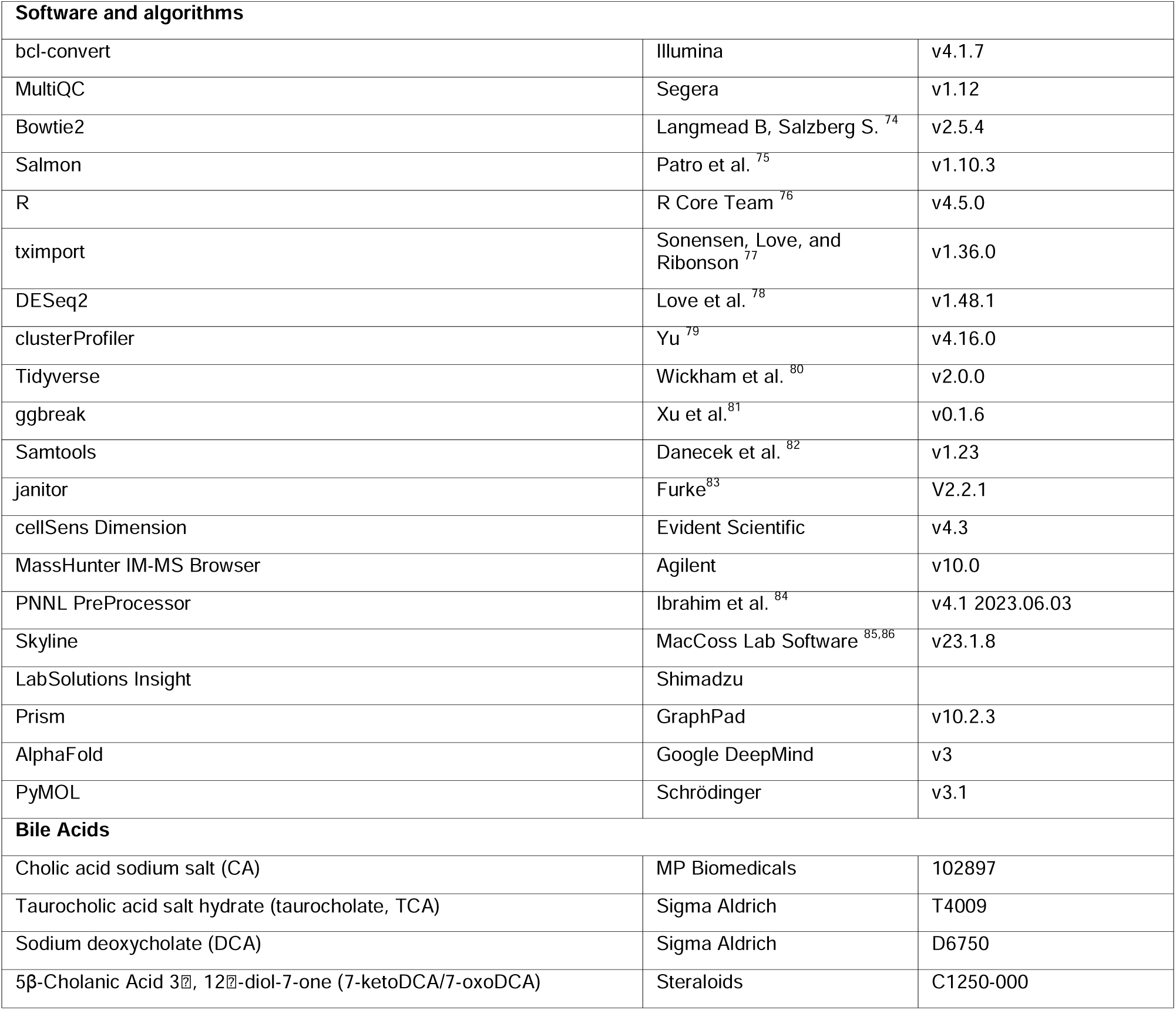

### EXPERIMENTAL MODELS

#### Strains and growth conditions

These experiments used the clinically relevant strain of *C. difficile*, R20291. *C. difficile* spores were germinated from stocks maintained at 4°C on brain heart infusion (BHI) agar (VWR, 90003-258) supplemented with 100 mg/L L-cysteine (Alfa Aesar, A10435) and 10% w/v taurocholate (Sigma Aldrich, T4009), referred to as TBHI. Additionally, the commensal strains *C. scindens* VPI 12708, and *C. hiranonis* TO 931 were used. *C. scindens* and *C. hiranonis* were stored at -80°C in 30% glycerol. *C. scindens* was grown in BHI broth (VWR, 90000-060) supplemented with 100 mg/L L-cysteine *C. hiranonis* was grown in BHI supplemented with 100 mg/L L-cysteine and 2 μM hemin (Alfa Aesar, A11165). *C. difficile* was grown in BHI supplemented with 100 mg/L L-cysteine for all spent media and competition studies with *C. scindens*. Cultures of *C. difficile* were supplemented with 2 μM hemin in addition to L-cysteine for spent media and competition studies with *C. hiranonis*. All cultures were grown in anaerobic conditions with 2.5% hydrogen (Coy, USA) at 37°C. *C. difficile* and *C. scindens* were maintained and enumerated on BHI agarsupplemented with 100 mg/L L-cysteine, and *C. hiranonis* was maintained and enumerated on BHI agar supplemented with 100 mg/L L-cysteine and 2 μM hemin. For spent media and competition studies with *C. hiranonis*, *C. difficile* was also enumerated and vegetative cells were maintained on BHI agar supplemented with 100 mg/L L-cysteine and 2 μM hemin. *C. difficile* spores were heated at 65°C for 20 min and enumerated on TBHI agar for *in vivo* and *in vitro* experiments. All *in vivo C. difficile* samples were enumerated on BHI agar.

#### Animals and housing

C57BL/6J mice (male and female, 4-9 weeks old) purchased from the North Carolina State University Gnotobiotic Animal Core were used for the experimental infections. Mice were housed in a sterilized biosafety cabinet with autoclaved bedding, water and food. Cage changes were performed weekly within the sterilized biosafety cabinet. All mice were subjected to a 12-hour light and 12-hour dark cycle. Animal experiments were conducted in the Laboratory Animal Facilities located on the NCSU CVM campus. The animal facilities are equipped with a full-time animal care staff coordinated by the Laboratory Animal Resources (LAR) division at NCSU. The NCSU CVM is accredited by the Association for the Assessment and Accreditation of Laboratory Animal Care International (AAALAC). Trained animal handlers in the facility fed and assessed the status of animals several times per day. Those assessed as moribund were humanely euthanized by CO_2_ asphyxiation. This protocol is approved by NC State’s Institutional Animal Care and Use Committee (IACUC).

#### Gnotobiotic mouse model of *C. difficile* infection

Groups of 4–9-week-old, male and female C57BL/6J mice (n= 83) were entered into a sterilized biosafety cabinet and had stool test culture negative for bacteria for quality control. Mice were then randomly sorted into the following experimental groups: (1) monocolonized commensal only; (2) CDI only; (3) commensal and CDI. On day -7 all mice with the exception of those in the CDI only treatment group were inoculated via oral gavage with ∼10^7^ CFU vegetative cells of either *C. scindens* VPI 12708 or *C. hiranonis* TO 931 as the commensal Clostridia. Mice were then allowed to obtain stable colonization of their respective commensal, monitored by bacterial enumeration of feces on BHI agar. On day 0 all mice, with the exception of those in the monocolonized commensal only treatment group, were challenged via oral gavage with ∼10^5^ *C. difficile* R20291 spores. Progression of *C. difficile* infection (CDI) was monitored closely starting on day 0 for up to 2 days post challenge. Animals were humanely euthanized by CO_2_ asphyxiation followed by cervical dislocation prior to necropsy. During necropsy content and tissue snips were taken from the cecum and flash frozen in liquid nitrogen then stored at -80^0^C until further analysis. Some of the cecal content that was collected was used for bacterial enumeration on the day of necropsy.

#### Challenging gnotobiotic mice with *C. scindens* and *C. hiranonis* BSH

Germ-free mice were administered 10 µg *C. hiranonis* BSH by oral gavage once daily on day -1, day 0, and day 1. BSH solution was prepared by dialyzing into PBS, pH 7.4. Activity of the enzyme was confirmed before and after oral gavage via the ninhydrin assay.

### METHOD DETAILS

#### Competition experiments

Competition experiments were performed following previously described methods ^41,44^ (Reed et al., 2020). Briefly, overnight cultures of *C. difficile* and either *C. scindens* or *C. hiranonis* were prepared and incubated in anaerobic conditions at 37°C. After 16 hr of incubation, *C. difficile* cultures were back diluted 1:10 and 1:5 in fresh BHI and incubated to allow doubling, approximately 4 hr. Commensal cultures, either *C. scindens* or *C. hiranonis*, were also back diluted 1:5 and 1:10 and allowed to double for 4 hr. Culture doubling was determined by measuring OD_600_ and comparing to the starting OD_600_ value. Doubled cultures were then used to prepare the competition assay. For *C. scindens* competition experiments, both *C. difficile* and *C. scindens* cultures were diluted to a starting concentration of ∼1 x 10^5^ CFU/mL. To prepare the final starting cultures, each strain was added to fresh BHI broth at a 1:1 ratio in a final volume of 10 mL. For cultures where BAs were also added, volumes of broth added were adjusted to reach a final concentration of 1.25 mM CA (MP Biomedicals, 102897) or 1.25 mM DCA (Sigma Aldrich, D6750). Starting cultures were enumerated via spread-plating before being incubated for 24 hr in anaerobic conditions at 37°C. After incubation, cultures were again enumerated at the 24 hr timepoint via spread plating. Additionally, 500 μL of culture was reserved for later toxin assays and 500 μL of culture was reserved for targeted BA metabolomics. The remaining 9 mL of culture was quenched in a 1:1 mix of ethanol and acetone for later RNA isolation. The same protocol was followed for *C. hiranonis* with the following modifications: all cultures were grown in BHI supplemented with hemin, as described above, and starting concentrations of *C. difficile* and *C. hiranonis* were set to ∼ 1 x 10^6^ CFU/mL to account for *C. hiranonis* growth defects at a lower starting density.

#### Spent media experiments

Spent media experiments were performed as previously described ^41^, with the addition of a toxin assay on samples collected after 24 hr of incubation. Overnight cultures of either *C. scindens* or *C. hiranonis* were prepared by inoculating fresh BHI broth, supplemented with hemin for *C. hiranonis* cultures, with either commensal and allowed to grow overnight. Cultures containing bile acids were prepared the same way, with the addition of 2.5 mM CA or DCA. At approximately 19 hr post inoculation, cultures were filter sterilized using a 0.22 μm PVDF syringe-driven filter (Genesee Scientific, 25-239). Simultaneously, overnight cultures of *C. difficile* were prepared in fresh BHI and incubated overnight in anaerobic conditions at 37°C. After 16 hr of growth, the overnight cultures were back-diluted 1:10 and 1:5 in fresh BHI and allowed to double for 4 hr. Doubling was confirmed by comparing the OD_600_ value measured after incubation to the starting OD_600_ value, and cultures were prepared at a final volume of 5 mL in the following ratio: 1 mL of fresh BHI with *C. difficile* at ∼ 1 x 10^6^ CFU/mL and 4 mL of filter sterilized spent media. Cultures were enumerated at 0 hr by drip plating to ensure a consistent starting inoculum before incubating at 37°C for 24 hr. At 24 hr post inoculation, cultures were again enumerated by drip-plating. Samples were also collected for later toxin assays and targeted bile acid metabolomics, with 500 μL being reserved for each assay (1 mL total reserved). The remaining 4 mL of culture was quenched in a 1:1 mix of ethanol and acetone for later RNA isolation.

#### Bile acid supplementation experiments

Bile acid supplementation experiments were done by growing *C. difficile* alone or with 0.625 mM of either CA, DCA, or 7-oxoDCA (Steraloids, C1250-000). Cultures were also grown in spent media from *C. hiranonis* +/- 2.5 mM CA, which served as a baseline. An ethanol only control was also prepared to account for the effects of the vehicle for 7-oxoDCA. All cultures were prepared at a final volume of 10 mL. Cultures were incubated at 37°C and samples were collected for spent media sporulation experiments and Vero cell cytotoxicity assays described below.

#### Spent media sporulation experiments

Sporulation was quantified from samples prepared following the above BA supplementation preparation method. At 24, 48, and 72 hr post inoculation, a 500 μL aliquot was taken from each culture and split into three separate samples: 100 μL for enumeration, 200 μL for later toxin assays, and 200 μL for archival purposes. From the enumeration aliquot, 20 μL was used for drip plating on BHI to quantify vegetative cells. The aliquot was then passed out of the anaerobic chamber and heated at 65°C for 20 min to kill vegetative cells, leaving only spores. The heated aliquot was passed back into the anaerobic chamber and used for drip plating on TBHI to quantify spores.

#### Vero cell cytotoxicity assay

Toxin activity was measured using a Vero cell cytotoxicity assay. Vero cells are grown and maintained in DMEM media (Gibco Laboratories, 11965-092) with 10% fetal bovine serum (Gibco Laboratories, 16140-071) and 1% penicillin-streptomycin solution (Gibco Laboratories, 15070-063). Cells were incubated with 0.25% trypsin (Gibco Laboratories, 25200-056), washed with 1x DMEM media, and harvested by centrifugation at 250 x g for 5 min. Cells were seeded at 1 x 10^4^ cells per well in a 96-well flat-bottom microtiter plate and incubated overnight at 37°C/5% CO_2_. The fecal pellets or cecal content was diluted (1:10) with PBS and filtered before testing, and the spent media samples were only filtered before testing. The samples were further diluted 10-fold to a maximum of 10^-6^. Sample dilutions were incubated 1:1 with PBS (for all dilutions) or antitoxin (TechLab, T5003) for 40 min at room temperature. Following the incubation, these mixtures were added to the Vero cells and plates were incubated overnight at 37°C/5% CO_2_. Vero cells were viewed under 200X magnification for rounding after overnight incubation. The cytotoxic titer was defined as the reciprocal of the highest dilution that produced rounding in 80% of Vero cells for each sample. Vero cells treated with purified *C. difficile* toxin A were used as controls. For *in vitro* samples, the limit of detection for this assay was a titer of 1. *In vivo* samples had a limit of detection of 2 for this assay due to the initial dilution of the fecal pellet or cecal content.

#### RNA isolation from *in vivo* samples

RNA was isolated from cecal content collected from mice on day 2 post *C. difficile* challenge. Approximately 200 mg of frozen cecal content was aliquoted into a new 1.5 mL microcentrifuge tube and 800 μL 1x DNA/RNA Shield (Zymo, R1100) was added to each sample and pipette mixed with a wide-bore P1000 tip (Genesee Scientific, 22-429P). 600 μL of the resuspended sample was added to Zymo ZR BashingBead Lysis Tubes 0.1mm (S6012-50) pre-filled with an equal volume of Trizol (Invitrogen, 15596026). Bead lysis tubes were secured in a vortex mixer with bead tube adapter and run for 10 minutes at maximum speed. Tubes were then centrifuged at 12,000xg for 10 min at 4°C. After centrifugation, the upper aqueous phase from each sample was added to a fresh 1.5 mL microcentrifuge tube containing 750 μL Trizol and pipette mixed. Sample tubes were incubated for 15 min, before 200 μL of chloroform was added and the tubes were inverted quickly for 20 sec. These tubes were then incubated for an additional 15 min before being centrifuged at 12,000xg for 15 min at 4°C. The clear aqueous phase was removed after centrifugation and transferred into a new 1.5 mL microcentrifuge tube. An equal volume of pure molecular grade 200 proof ethanol was added to each sample, mixed well, and used as input for the Zymo RNA Clean and Concentrator kit (Zymo, R1017). Resulting RNA was quantitated by Qubit RNA High Sensitivity Assay (Invitrogen, Q32852). RNA integrity was measured with TapeStation High Sensitivity RNA ScreenTape (Agilent, 5067-5579) at the North Carolina State University Genomic Sciences Laboratory.

#### RNAseq of *in vivo* samples

RNA isolated from mouse cecal content was sequenced at the Roy J. Carver Biotechnology Center at the University of Illinois at Urbana-Champaign. Purified DNAsed total RNAs were run on a Fragment Analyzer (Agilent, CA) to evaluate RNA integrity. The total RNAs were converted into individually barcoded polyadenylated mRNAseq libraries with the Kapa Hyper Stranded mRNA library kit (Roche, 08098123702), with prior removal of rRNAs with the FastSelect HMR and Bacteria kits (Qiagen, 335925). Libraries were barcoded with Unique Dual Indexes (UDI’s) which have been developed to prevent index switching. The final libraries were quantitated with Qubit (ThermoFisher, MA) and the average cDNA fragment sizes were determined on a Fragment Analyzer. The libraries were diluted to 10 nM and further quantitated by qPCR on a CFX Connect Real-Time qPCR system (Biorad, CA) for accurate pooling of barcoded libraries and maximization of number of clusters in the flowcell. The barcoded RNAseq libraries were loaded on one 10B lane on a NovaSeq X Plus (Illumina) for cluster formation and sequencing. The libraries were sequenced as from both ends of the fragments for a total of 150bp from each end.

#### RNAseq analysis of *in vivo* samples

Resulting FASTQ files were demultiplexed with the bcl-convert v4.1.7 Conversion Software (Illumina). Adaptors were trimmed from the 3’ end of reads, and quality control for each output sequence was performed with MultiQC v1.12 (Seqera). For each sample, forward and reverse host reads were mapped with Bowtie2 (v2.5.4). Unmapped reads were isolated and split into forward and reverse reads with Samtools (v1.23). Resulting unmerged paired-end reads were mapped to the *C. difficile* R20291 genome (NCBI RefSeq NC_013316.1), or the *C. hiranonis* TO-931 genome (NCBI RefSeq NZ_CP036523) using Salmon (v1.10.3). For reads from mice cocolonized with both *C. difficile* and *C. hiranonis*, reads were mapped to both reference genomes. Resulting mapped reads were imported into R (v4.5.0) with tximport (v1.36.0).. Differential expression analysis was done to compare either *C. hiranonis* or *C. difficile* between monocolonized and cocolonized mice with DESeq2 (v1.48.1). Resulting data was visualized using ggplot2 in the tidyverse package suite (v2.0.0). Axis breaks were added by ggbreak (v0.1.6). All data transformations were done with tidyverse packages (v2.0.0). Significantly differentially expressed genes associated with *C. difficile* sporulation were manually annotated by first aligning to a list of previously identified essential sporulation-associated genes ^45^. This output was then used to search UniProt for further specific annotations. Individual annotations were also made from relevant literature sources if previous works had identified a particular gene’s function ^87,88^. Non-sporulation associated genes were classified by KEGG pathway using clusterProfiler (v4.16.0).

#### RNA isolation from *in vitro* samples

RNA was isolated from bacterial cultures as previously described ^69^. Briefly, cultures were fixed with an equal volume of a 1:1 mixture of ethanol and acetone and frozen at -80°C until extraction. Before extraction, frozen cultures were thawed and spun at 4,000 rpm for 10 minutes at 4°C and the supernatant was decanted. The cell pellet was resuspended in a 1:100 solution of water and β-mercaptoethanol and transferred to a fresh 1.5 mL microcentrifuge tube. The resuspended cell pellet was centrifuged at 16,000xg for one minute at room temperature. After centrifugation the resulting supernatant was decanted, and the cell pellet was resuspended in 1 mL cold Trizol (Invitrogen, 15596026) and allowed to sit at room temperature for 15 min with occasional inversion. 200 μL of chloroform was then added to each tube before vigorous inversion for 20 sec. Tubes were allowed to sit for an additional 15 min before being centrifuged at 16,000xg for 15 min at 4°C. Centrifugation caused phase separation, yielding approximately 500 μL of clear aqueous phase that was transferred to a fresh 1.5 mL microcentrifuge tube. The aqueous phase was mixed with an equal volume of 100% ethanol and used as input for extraction following the Direct-zol RNA Miniprep kit protocol (Zymo, R2050). On-column DNaseI treatment was used according to kit protocol. Isolated RNA was quantitated via Qubit High Sensitivity RNA assay (Invitrogen, Q32852) and RNA integrity was measured with TapeStation High Sensitivity RNA ScreenTape (Agilent, 5067-5579) at the North Carolina State University Genomic Sciences Laboratory. Remaining RNA was frozen at -80°C.

#### RNAseq of *in vitro* samples

RNA isolated from *in vitro* bacterial cultures was sequenced at the Roy J. Carver Biotechnology Center at the University of Illinois at Urbana-Champaign. Purified DNAsed total RNAs were run on a Fragment Analyzer (Agilent, CA) to evaluate RNA integrity. The total RNAs were converted into individually barcoded polyadenylated mRNAseq libraries with the Watchmaker mRNA library kit (Watchmaker, 7BK0001), with prior removal of rRNAs with FastSelect HMW +23/16S probes (Qiagen, 335925). Libraries were barcoded with Unique Dual Indexes (UDI’s) which have been developed to prevent index switching. The final libraries were quantitated with Qubit (ThermoFisher, MA) and the average cDNA fragment sizes were determined on a Fragment Analyzer. The libraries were diluted to 10nM and further quantitated by qPCR on a CFX Connect Real-Time qPCR system (Biorad, CA) for accurate pooling of barcoded libraries and maximization of number of clusters in the flowcell. The barcoded RNAseq libraries were loaded on one 10B lane on a NovaSeq X Plus for cluster formation and sequencing. The libraries were sequenced as from both ends of the fragments for a total of 150bp from each end.

#### RNAseq analysis of *in vitro* samples

Resulting FASTQ files were demultiplexed with the bcl-convert v4.1.7 Conversion Software (Illumina). Adaptors were trimmed from the 3’ end of reads, and quality control for each output sequence was performed with MultiQC v1.12 (Seqera). Paired-end reads were mapped to the *C. difficile* R20291 genome (NCBI RefSeq NC_013316.1) using Salmon (v1.10.3) and mapped reads were imported into R (v4.5.0) with tximport (v1.36.0). Differential expression analysis was performed with DESeq2 (v1.48.1). All necessary data transformations were done with tidyverse packages (v2.0.0) and data cleaning was performed with janitor (v2.2.1). Genes were classified by KEGG Pathway with clusterProfiler (v4.16.0) and resulting data was visualized using ggplot2 within the tidyverse suite (v2.0.0).

#### Histopathological examination of the mouse cecum and colon

The cecum and colon were prepared for histology, at the time of necropsy, by placing representative sections into tissue cassettes, which were stored in 10% neutral buffered formalin for 72 h at room temperature before being transferred to 70% ethanol at room temperature for long-term storage. Tissue cassettes were processed, paraffin751 embedded, and sectioned at 4 um thickness, and hematoxylin and eosin stained for histopathological examination (University of North Carolina Animal Histopathology & Lab Medicine core). Histological specimens were randomized and scored in a blinded manner by a board-certified veterinary pathologist (ER). Edema, leukocyte infiltration, and epithelial damage were each scored 0-4 for both cecum and colon based on a previously published numerical scoring scheme. Edema scores were as follows: 0, no edema; (1) mild edema with minimal (up to 2Å∼) multifocal submucosal expansion or a single focus of moderate (2–3Å∼) submucosal expansion; (2) moderate edema with moderate (2–3Å∼) multifocal submucosal expansion; (3) severe edema with severe (3Å∼) multifocal submucosal expansion; (4) same as score 3 with diffuse submucosal expansion. Cellular infiltration scores were as follows: 0, no inflammation; (1) minimal multifocal neutrophilic inflammation of scattered cells that do not form clusters; (2) moderate multifocal neutrophilic inflammation (greater submucosal involvement); (3) severe multifocal to coalescing neutrophilic inflammation (greater submucosal ± mural involvement); (4) same as score 3 with abscesses or extensive mural involvement. Epithelial damage was scored as follows: 0, no epithelial changes; (1) minimal multifocal superficial epithelial damage (vacuolation, apoptotic figures, villus tip attenuation/necrosis); (2) moderate multifocal superficial epithelial damage (vacuolation, apoptotic figures, villus tip attenuation/necrosis); (3) severe multifocal epithelial damage (same as above) +/− pseudomembrane (intraluminal neutrophils, sloughed epithelium in a fibrinous matrix); (4) same as score 3 with significant pseudomembrane or epithelial ulceration (focal complete loss of epithelium). Photomicrographs were captured on an Olympus BX43 light microscope with a DP27 camera using the cellSens Dimension software. The scale bar in each histopathology picture measures 100 μm in size.

#### BA metaboloimcs extraction method for mouse cecal content

Cecal samples were processed using a bile acid extraction protocol with minor modifications, as previously described^24^. Approximately 150 mg of cecal content per sample was aliquoted into 2 mL Omni microtubes containing 2.38 mm metal beads (Omni International, Kennesaw, GA, USA; catalog no. 19-620) for homogenization. The pre-extraction solvent system consisted of an 85:15 (v/v) organic:aqueous solution, prepared by combining equal volumes of LC-MS grade acetonitrile and methanol (1:1, v/v); (Fisher Scientific, Waltham, MA) with 20 mM potassium phosphate buffer. This resulted in a final extraction solvent composition of 50:50 (v/v) acetonitrile/methanol containing 3 mM phosphate. A 50 µM ^13^C-labeled internal standard (Cambridge Isotope Laboratories, Tewksbury, MA, MSK-BA1 and MSK-BA-2) mixture was added to the solvent at a 1:100 (v/v) ratio, achieving a final internal standard concentration of 0.5 µM in the pre-extraction buffer.

Samples were normalized by mass, adjusting extraction volumes to a 1:8 (mg:µL) ratio according to the precise weight of each individual sample, as detailed in Supplemental Table S1. Homogenization was performed using a Fisherbrand™ Bead Mill 24 Homogenizer (Fisher Scientific, Hampton, NH, catalog no. 15-340-163), ensuring complete tissue disruption. Following homogenization, samples were centrifuged at 10,000 × g for 15 min at 4°C using an Eppendorf 5810R centrifuge to pellet insoluble debris. From the saved supernatant which was transferred into a clean 2 mL Thermo microcentrifuge tube,100 µL was used and mixed with 100 µL methanol (Fisher Scientific, catalog no. 67-56-1). Samples were vortexed briefly, then shaken at 600 rpm for 20 min at room temperature using a Thermo MixMate shaker (Hamburg, Germany, 5353000529). Prior to LC-IMS-MS analysis, extracts were filtered through a 0.2 µm PTFE membrane spin filter (Millipore Sigma, Darmstadt, Germany, UFC30LG25) by centrifugation at 10,000 × g for 1 min. Filtrates were transferred to Agilent LC vials (Agilent, Santa Clara, CA, catalog no. 5188-6591) and stored at −80°C until analysis.

#### Extraction method for germ-free mouse cecal content

Extraction of germ-free mouse cecal samples followed the same procedure described above for *in vivo* samples. To maintain consistency, germ-free samples were extracted using the same pre-prepared extraction buffer as utilized for the *in vivo* sample procedure. However, it should be noted that germ-free samples were processed in different extraction batches, using distinct internal standard batches during the pre-extraction step. Variability due to internal standard batch differences and homogenization efficiency during the bead mill procedure was not controlled between the two groups. Consequently, absolute internal standard intensities observed in germ-free samples are not directly comparable to those from the *in vivo* cohort.

#### Extraction method for bacterial cultures

A ^13^C-labeled internal standard mixture described above was incorporated into LC-MS-grade methanol extraction buffer (Fisher Scientific, Waltham, MA) at a 1:100 (v/v) dilution, resulting in a final internal standard concentration of 0.5 µM. For each bacterial culture sample, 100 µL of culture was transferred into a 1.5 mL microcentrifuge tube and mixed with 400 µL of internal standard-containing methanol buffer. Samples were vortexed at 2000 rpm for 5 min using a Thermo MixMate shaker (Hamburg, Germany; catalog no. 5353000529) followed by sonication on ice for 15 min. Extracts were incubated overnight at −20°C. Following incubation, samples were centrifuged at 13,000 × g for 10 min at 4°C using an Eppendorf 5424R centrifuge. Then, 300 µL aliquots of the resulting supernatants were transferred into fresh 1.5 mL microcentrifuge tubes and stored at −80°C until LC-IMS-MS analysis.

#### LC-IMS-MS analysis of *in vivo* samples

All extracted bile acid samples were analyzed using a liquid chromatography–ion mobility spectrometry–mass spectrometry (LC-IMS-MS) platform consisting of an Agilent 1290 Infinity II UHPLC coupled to a 6560 Ion Mobility QTOF mass spectrometer (Agilent Technologies, Santa Clara, CA, USA) ^89,90^. Instrument calibration was performed prior to analysis using the Agilent ESI-L Low Concentration Tuning Mix (catalog no. G1969-85000) to ensure accurate mass measurements and precise collision cross-section (CCS) determination. Chromatographic separation was performed on a Restek Raptor C18 column (1.8 μm, 2.1 × 50 mm; catalog no. 9304252) equipped with a ZORBAX Eclipse Plus C18 Guard Column (1.8 µm, 2.1 x 5 mm, catalog no. 823750-901), both maintained at 60°C under neutral mobile-phase conditions. The mobile-phase system consisted of 5 mM ammonium acetate in water (mobile phase A) and methanol/acetonitrile (50:50, v/v; mobile phase B). Samples (6 µL) were injected and separated at a constant flow rate of 0.5 mL/min according to the gradient detailed in Supplemental Table S2. Ion mobility separations were conducted in a 78 cm drift tube utilizing nitrogen as the buffer gas at 3.95 Torr. Data were acquired in negative ion mode using 4-bit multiplexing over the mass range of 50–1700 m/z. Drift times were converted to CCS values using the standard tuning mix, and ion transmission parameters were optimized to enhance peak capacity. Trap fill and release times were set at 3.9 ms and 150 µs, respectively. Further details of MS parameters are provided in Supplemental Table S3. Raw data files were processed using Agilent MassHunter IM-MS Browser (v10.0), demultiplexed using demultiplexed using a PNNL PreProcessor (v4.1 2023.06.03), and the resulting .DeMP.d files were imported into Skyline software ^84–86^ (v23.1.8; MacCoss Lab Software, Seattle, WA) for targeted data extraction. Skyline’s ion mobility filtering feature, employing experimentally derived arrival times, was utilized to enhance peak detection specificity. Peak quantification was performed using area-under-the-curve (AUC) metrics, and features with mass errors exceeding ±10 ppm and AUC below a defined threshold of 1000 were excluded. A comprehensive list of bile acid targets, including retention times, *m/z* values, and calculated CCS values, is provided in Supplemental Table 4.

Resulting bile acid AUC data was used as input for principal component analysis using the stats package in base R (v4.5.0). Comparisons between conditions were also made using the t-test function from the stats package, and resulting data was visualized with ggplot2 within the tidyverse suite (v2.0.0). All necessary data manipulations were performed with tidyverse packages (v2.0.0) and data cleaning was performed with janitor (v2.2.1).

#### Differential scanning fluorometry

DSF was performed in a similar manner as described previously ^91^. TcdB protein was diluted to 0.05 mg/mL in phosphate buffer (100mM KPO4, 150mM NaCl, pH7) containing 5X SYPRO Orange (ThermoFisher, S6650) and a serial dilution of test compound in 1% final DMSO. A Biorad CFX96 qRT-PCR thermocycler was used to establish a temperature gradient from 42°C to 65°C in 0.5-degree increments, at 5 seconds per increment, while simultaneously recording the increase in SYPRO Orange fluorescence as a consequence of binding to hydrophobic regions exposed on unfolded proteins. The Bio-Rad CFX Manager 3.1 software was used to integrate the fluorescence curves in order to calculate the melting points, which were then plotted using Prism software (Graphpad, v10.2.3).

#### Cell rounding assay

IMR-90 cells were purchased from ATCC. Mycoplasma monitoring was performed using the MycoAlert Plus kit (Lonza, LT07), and Mycostrip Mycoplasma Detection Kit (Invivogen, rep-mys); in all cases the IMR90 cells tested negative for mycoplasma contamination. IMR-90 cells were grown in EMEM (Wisent, 320-005-CL) supplemented with 10% FBS (Wisent, 090-450) and penicillin-streptomycin (complete EMEM) and were seeded in 96well Cellbind plates (Corning) at a density of 8,000-10,000 cells/well. The next day, the media was exchanged with serum free EMEM (SFM) containing 1 μM Celltracker Orange CMRA (ThermoFisher, C34551). After 60 minutes, excess dye was removed by media exchange with 90 μL SFM per well. An Agilent Bravo liquid handler was used to deliver 0.4 μL of serially diluted compound from the source plate to the cell plate, immediately followed by 10 μL of 5 pM TcdB (diluted in SFM), representing a final concentration of 0.5 pM toxin, previously established as ∼EC99 levels of cytopathology. The cell plates were returned to the incubator for 2.5hrs before imaging. Celltracker-labelled cells were evaluated on a Cellomics ArrayScan VTI HCS reader (ThermoFisher) using the Target Acquisition mode, a 10X objective and a sample rate of 150 objects per well. After recording all image data, the cell rounding and shrinking effects of TcdB intoxication were calculated using the cell rounding index (CRI), a combined measure of the length to width ratio (LWR) and area parameters. The % inhibition was calculated as the ratio between the sample well and the average toxin-untreated controls after subtracting the average vehicle-only control values and plotted using Prism software (GraphPad, v10.2.3).

#### Cloning of *C. hiranonis bsh* and recombinant protein expression

The nucleotide sequence for *C. hiranonis bsh* was codon optimized and synthesized as a gene fragment (Integrated DNA Technologies). The *bsh* gene was cloned into the pET-28b(+) vector (Sigma Aldrich, 69865-M) using the NEBuilder HiFi DNA Assembly Kit (New England Biolabs, E5520S), and proper insertion of the gene was confirmed via sequencing (Eton Bioscience). *C. hiranonis bsh* plasmid DNA was transformed into *E. coli* BL21 (DE3)pLysS competent cells (ThermoFisher Scientific, C606010). Overnight starter cultures were prepared by inoculating 5 mL of Luria-Bertani (LB) media (Sigma Aldrich, L3022) supplemented with kanamycin (30 µg/mL) (Sigma Aldrich, 60615) and chloramphenicol (34 µg/mL) (Sigma Aldrich, C0378) with an isolated colony of transformed BL21 (DE3)pLysS cells and grown overnight at 37°C with agitation at 225 RPM. The following day, the entire overnight culture was added to 1 L TB media (tryptone, yeast extract, glycerol, potassium phosphate) in a baffled 2 L flask, supplemented with kanamycin (30 µg/mL) (Sigma Aldrich, 60615), and grown at 37°C with agitation at 225 RPM. Once the OD_600_ reached ≈ 0.6-0.8, cultures were cooled before expression was induced with 0.5 mM isopropyl-β-d-thiogalactopyranoside (IPTG) (Sigma Aldrich, I6758). Cultures were grown overnight at 25°C and harvested by centrifugation at 4,000 x at 4°C for 20 min and stored at -80°C.

#### Recombinant protein purification

Cell pellets were resuspended (5 mL/g cells) in Lysis Buffer (50 mM sodium phosphate, 300 mM NaCl, 10 mM imidazole, 1 mg/mL lysozyme (Sigma Aldrich, SAE0152), 3 U/mL DNAse (ThermoFisher Scientific, AM2238), protease inhibitor tablet (ThermoFisher Scientific, S8830-20TAB), pH 7.5) and stirred for 30 min. The resuspended cells were lysed by sonication (Sonics VCX500) at 40% amplitude for 5 min with cycles of 5 s on, 10 s off. The lysate was clarified by centrifugation at 20,000 x g at 4°C for 20 minutes. The supernatant was applied to a column containing HisPur cobalt resin (ThermoFisher Scientific, PI89965) equilibrated in Buffer A (50 mM sodium phosphate, 300 mM NaCl, 10 mM imidazole, pH 7.5). Following collection of the flow-through, the column was washed with 20 mL Buffer A before protein was eluted with 10 mL Buffer B (50 mM sodium phosphate, 300 mM NaCl, 150 mM imidazole (Sigma Aldrich, I2399), pH 7.5. Fractions containing *C. hiranonis* BSH were assessed for purity using 10% acrylamide SDS-PAGE analysis (ThermoFIsher Scientific, XP00100BOX; Abcam, Ab119211). Protein concentration was measured using a Bradford assay (ThermoFisher Scientific, 23200). BSH fractions were pooled and dialyzed into Buffer C (50 mM sodium phosphate, 150 mM NaCl, 10% glycerol, 10 mM DTT, pH 7.5) before being flash frozen in liquid N_2_ and stored at -80°C.

#### BSH ninhydrin activity assay

Deconjugation of bile acids by the BSH was measured using a ninhydrin assay as described previously ^30,92^. In short, 50 µL reactions containing activity buffer (50 mM sodium phosphate, 10 mM DTT, pH 6.0), 9 mM bile acid, and 20 nM BSH were incubated at 37°C for 5 min before being quenched with 50 µL 15% (w/v) trichloroacetic acid. Standard solutions of 0-9 mM amino acid were also prepared. Following clarification of any formed precipitate, 25 µL of the quenched reaction/standard was added to 475 µL ninhydrin reaction buffer (300 µL glycerol, 125 µL 0.25% (w/v) ninhydrin solution, 50 µL 0.5 M sodium citrate, pH 5.5) and boiled at 90°C for 14 min. Color development was halted by placing solutions on ice, and two technical replicates for each condition were plated onto a 96-well plate. Absorbance was measured at 570 nm using a Tecan Infinite F200 Pro plate reader. Activity of the BSH, reported as v_0_/[E], was calculated as µM free amino acid released per second per µM BSH.

#### Extraction method for mouse cecal content for LC-MS/MS

Samples containing cecal content were prepared for LC-MS/MS at the McGuire VA Medical Center LC-MS Core Lab, Virgina Commonwealth University as previously described ^93^. Briefly, samples were resuspended with 50 µL methanol (MeOH)/acetonitrile (ACN) (1:1, v/v) and homogenized. The fecal homogenate (50 µL) was added to 950 µL of 1:1 MeOH/ACN and vortexed. Samples were then prepared by adding 20 µL fecal homogenate, 20 µL internal standard mixture, and 80 µL MeOH/ACN/H_2_O (5%/5%/90%, v/v/v) to a microcentrifuge tube, vortexed, and centrifuged at 16,000 x *g* for 5 min. All LC-MS/MS-grade solvents and chemicals were purchased from Fisher Scientific. Internal standard mixture contained *d*_4_-CA, *d*_4_-CDCA, *d*_4_-DCA, *d*_4_-LCA, *d*_4_-GCA, *d*_4_-GDCA, *d*_4_-GLCA, *d*_4_-TCA, *d*_4_-TCDCA, *d*_4_-TLCA, *d*_4_-CA-3-*S*, *d*_4_-LCA-3-*S* and *d*_4_-CDCA-3-*S* (Cayman Chemical Co., Ann Arbor, MI) in MeOH/ACN/H_2_O (20%/30%/50%, v/v/v) at a final concentration of 500 nM. Extracted sample was then injected for LC-MS/MS analysis at a volume of 3 μL. Add that this was done at VCU.

#### LC-MS/MS analysis of *in vivo* samples

LC analysis was performed using a Shimadzu CBM-20A CL communications bus module, equipped with two LC-30 AD CL pumps, a DGU-20A3R and a DGU-20A5A degassing unit, a SIL-30AC CL autosampler and a CTO-20A CL column-oven (40°C). Samples were injected onto a Thermo Scientific Hypersil Gold C18 column (100 x 2.1 mm; 1.9 µm particle size, 175 Å pore size). Separation was carried out using 0.1% (v/v) acetic acid in water (Mobile Phase A) and MeOH/ACN (1:1, v/v) (Mobile Phase B) at a flow rate of 0.2 mL/min. The gradient was held at 35% B for 2.5 min, increased linearly from 35% B to 85% B for 20.5 min (2.5-23 min), increased to 99% B for 5 min (23-28 min), then reequilibrated to 35% B for 2 min (28-30 min). MS analysis was performed using an LCMS-8060 CL triple quadrupole instrument with an electrospray ionization source (Shimadzu, Japan). The MS parameters were as follows: spray voltage; 3000V, heating block temperature; 400°C, nebulizing gas flow; 3 L/min, drying gas flow; 10 L/min, heating gas flow; 10 L/min, interface temperature 300°C, collision gas (argon) pressure; 1.3 mm Torr, collision energy; 11–85 eV, all in the negative ion mode. Data acquisition and analysis was performed using LabSolutions Insight (Shimadzu). Resulting bile acid abundances were used for statistical testing and graphing with GraphPad Prism (v10.2.3).

### QUANTIFICATION AND STATISTICAL ANALYSIS

All numbers of animals from *in vivo* experiments or biological replicates from *in vitro* experiments can be found in the associated figure legends and STAR Methods sections. Details on statistical testing and programs used for each analysis can be found in the associated figure legends and/or STAR Methods sections.

