## Supplementary figures and images for "Bile acid dependent attenuation of toxin mediated disease is independent of colonization resistance against *C. difficile*"

### Figures S1-S5

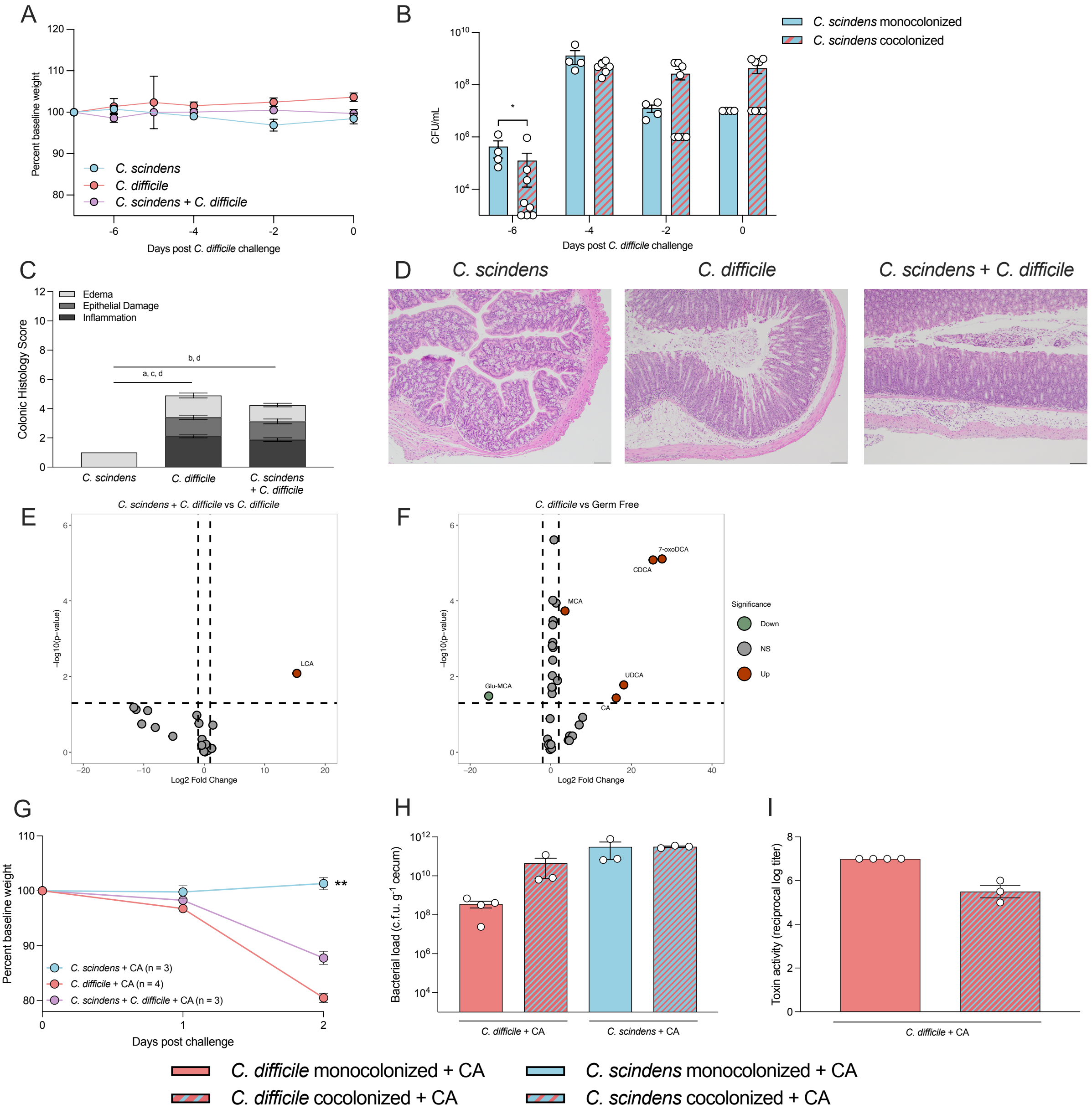

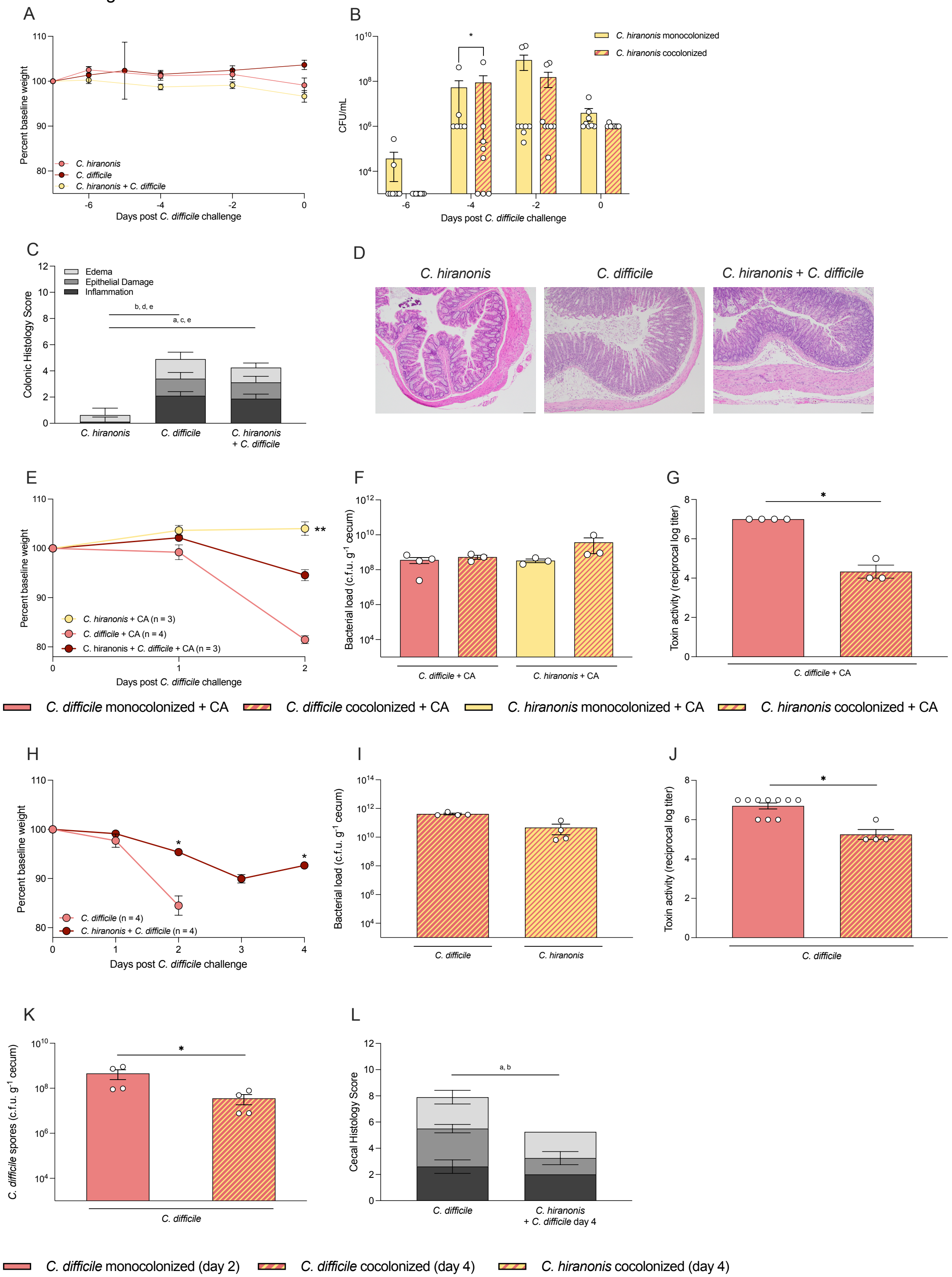

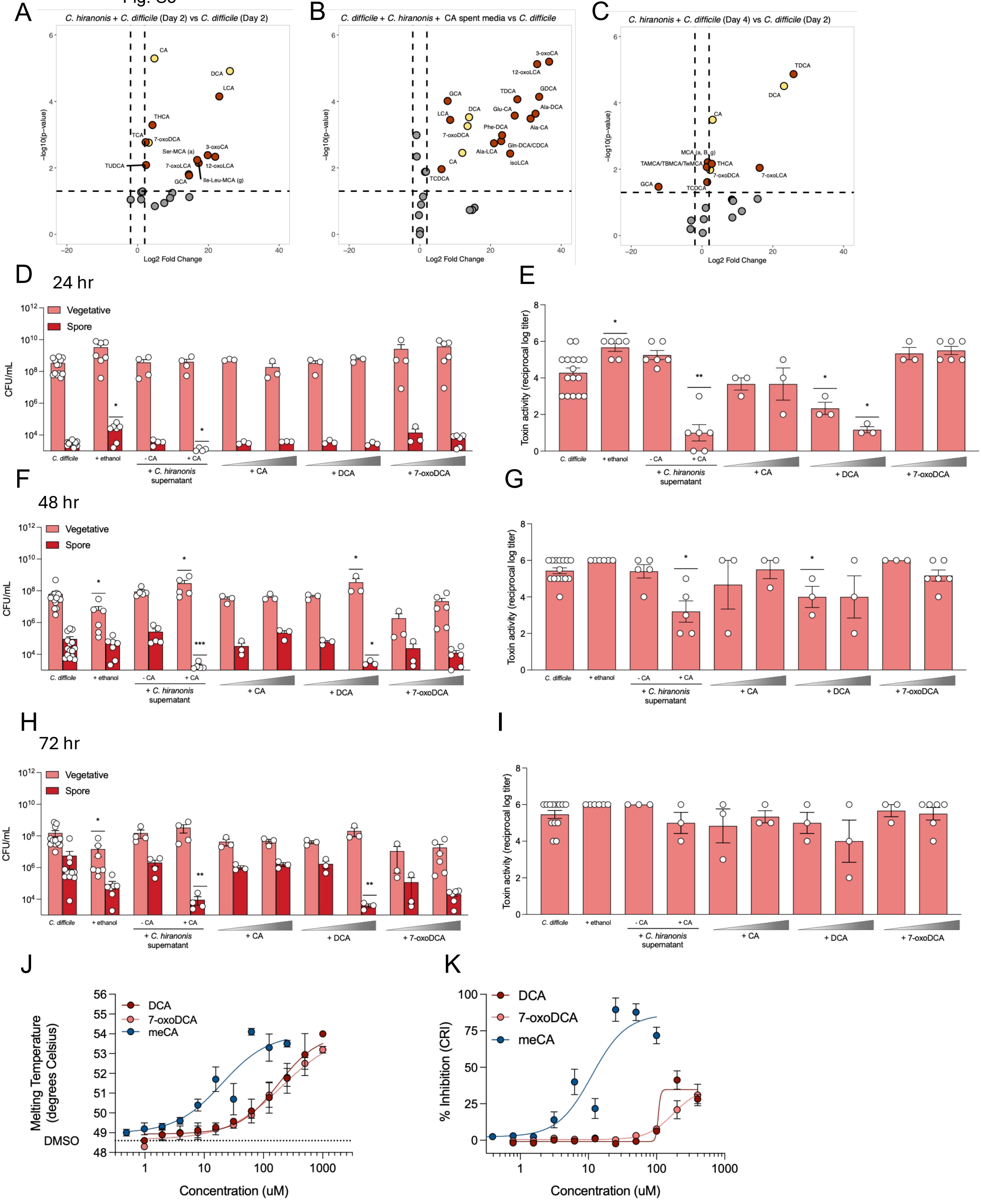

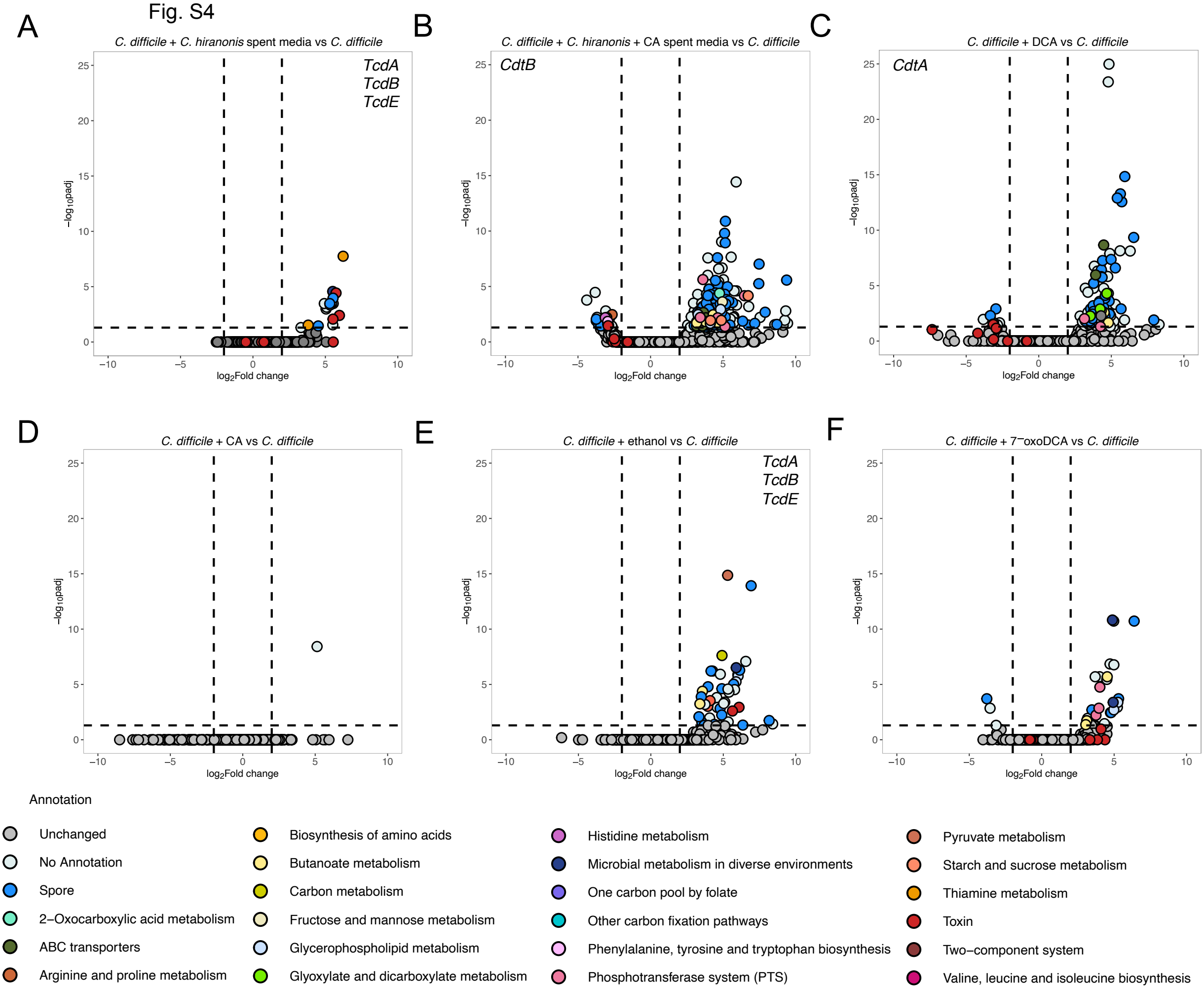

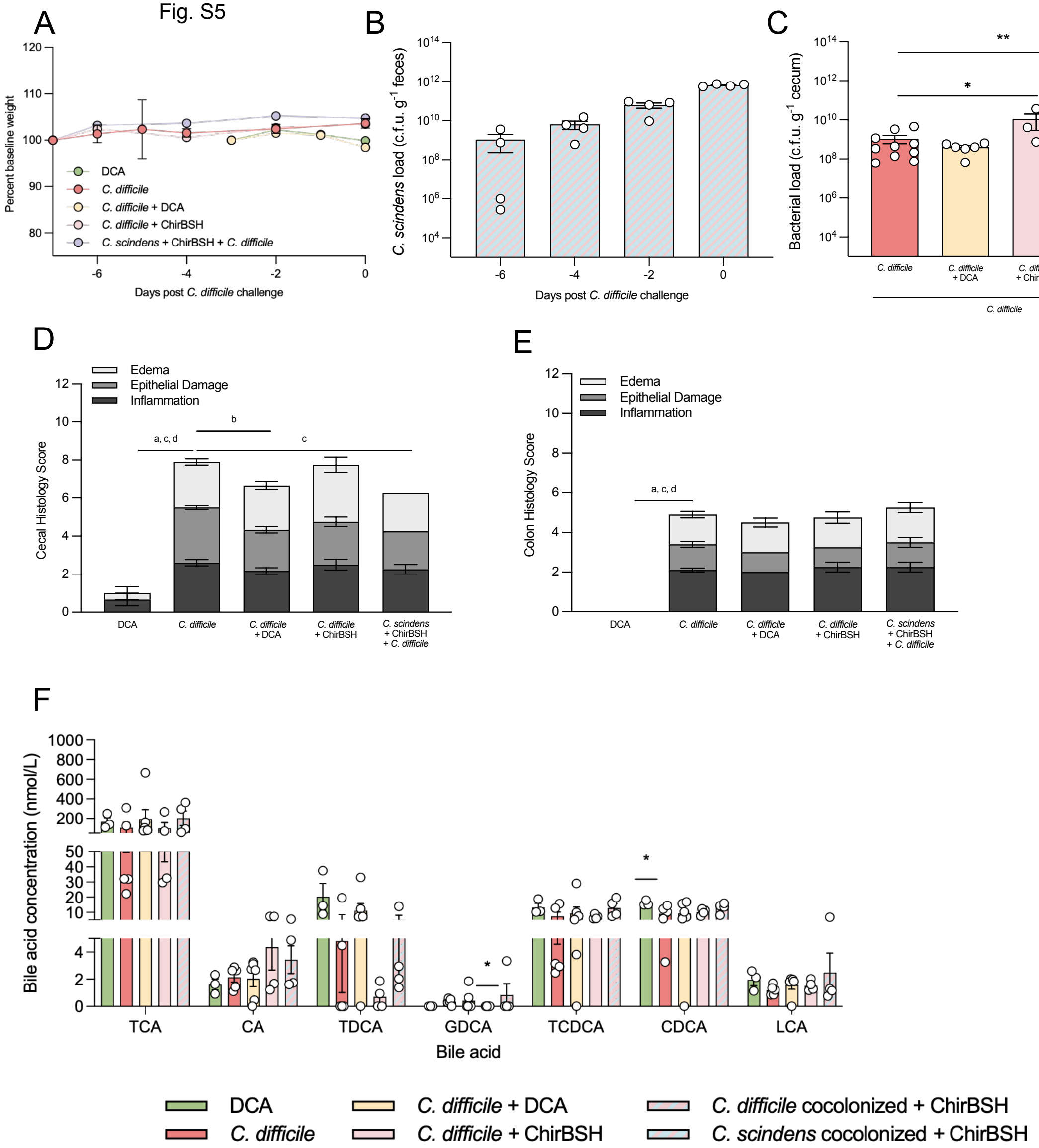
